# Longitudinal Phylogenetic Inference of Copy Number Alterations and Single Nucleotide Variants from Single-Cell Sequencing

**DOI:** 10.1101/2025.09.16.676596

**Authors:** Ethan Kulman, Quaid Morris, Rui Kuang

## Abstract

Longitudinal phylogenetic reconstruction reveals how cancers evolve over time and respond to treatments. Advances in targeted single-cell sequencing, combined with longitudinal sampling, now enable detailed longitudinal tracking of single nucleotide variants (SNVs) and copy number alterations (CNAs) at single-cell resolution. Here, we introduce LoPhy, the first method designed to reconstruct the evolution of SNVs and CNAs from these new longitudinal single-cell data. LoPhy is a sequential tree-building algorithm that reconstructs longitudinally-consistent phylogenies of SNVs and CNAs by maximizing a new factorized tree reconstruction objective. The algorithm incrementally grows a clone tree, adding SNVs and CNAs in the order they are observed across time points. Applied to a cohort of 15 acute myeloid leukemias (AMLs) and 4 TP53-mutated AMLs, LoPhy produced phylogenies that are biologically and temporally consistent with clinical observations, with many inferred CNAs validated by orthogonal bulk sequencing from the same cancer. These reconstructions highlight the role of CNAs in disease progression and resistance, revealing that AML clones selected after therapy are often defined by both large-scale CNAs and SNVs. More broadly, LoPhy can help uncover how SNVs and CNAs jointly shape the evolutionary trajectories of individual cancers at single-cell resolution. The LoPhy source code is available under a CC-BY-ND license at https://github.com/ethanumn/LoPhy.

**Author summary:** Longitudinal single-cell DNA sequencing is increasingly used to track somatic mutations in cancer, including single nucleotide variants (SNVs) and copy number alterations (CNAs). Phylogenetic analysis of such data can reveal the mutations that characterize key subpopulations of cancerous cells and how they evolve over time. However, no existing methods are designed to reconstruct the joint evolution of SNVs and CNAs from longitudinal single-cell data. To address this gap, we developed LoPhy, an algorithm that infers a phylogenetic tree from longitudinal single-cell data to capture the joint evolutionary history of SNVs and CNAs in an individual cancer. We applied LoPhy to simulated datasets and 19 acute myeloid leukemias (AMLs). LoPhy’s reconstructions reveal that AML subpopulations selected after treatment are frequently defined by both SNVs and large-scale CNAs—highlighting that joint longitudinal modeling of SNVs and CNAs is crucial for understanding disease progression and therapeutic resistance.

## Introduction

Cancer evolves through the accumulation of somatic mutations, most commonly single nucleotide variants (SNVs) and copy number alterations (CNAs) [1–3]. This evolutionary process gives rise to genetically distinct populations of cancer cells, known as *clones*, each defined by a unique genome. The clonal composition of a cancer can change over time due to selective pressures such as immune response or therapy. Longitudinal sequencing, which involves sampling and sequencing cells from the same cancer at multiple time points, enables researchers to track these evolutionary dynamics and gain insight into how cancers adapt and progress. Although the combined burden of SNVs and CNAs is known to influence cancer progression and therapeutic resistance [4–7], their joint effects over time within individual cells remain poorly understood. This gap largely reflects limitations of earlier single-cell DNA sequencing technologies, which could not reliably resolve both SNVs and CNAs in the same cells. Earlier approaches offered either broad genomic coverage with shallow depth (e.g., scWGS) or deep coverage of targeted regions with limited genomic breadth (e.g., targeted scDNA-seq). Recent advances in targeted single-cell DNA sequencing—particularly high-throughput microfluidic protocols known as *single-cell amplicon sequencing* —now enable the detection of both SNVs and CNAs [8, 9] at single-cell resolution. This technology has been applied to longitudinal cancer samples [8, 10], sequencing portions of oncogenes at *∼*40x per amplicon per cell.

While a few existing methods can reconstruct clone trees—phylogenetic trees describing the ancestral relationships among cancer clones—that capture the joint evolution of SNVs and CNAs from single-cell data [11–13], none are designed for longitudinal samples. Applying these methods to longitudinal samples either requires reconstructing clone trees separately for each time point—yielding disjoint evolutionary histories lacking temporal coherence—or pooling all samples together, which ignores sample-specific technical variation such as dropout rates and region-specific coverage. Although longitudinal phylogenetic reconstruction methods can exploit temporal structure and account for technical variation, existing approaches for single-cell data focus exclusively on either SNVs [14–16] or CNAs [17, 18], but not both.

Here, we introduce LoPhy, the first algorithm for reconstructing clone trees that capture the joint evolution of SNVs and CNAs from longitudinal single-cell amplicon sequencing data. The method relies on a factorized tree reconstruction objective—a novel formulation derived in this work. To optimize this objective, LoPhy uses a sequential tree-building strategy in which mutations are added to the tree in the order they are first detected across longitudinal samples, ensuring that mutations detected earlier in time are placed earlier in evolution. LoPhy generates a sequence of time-dependent trees, with each tree describing the evolutionary history up to a given time point and forming a subtree of all subsequent trees. Each tree is refined through stochastic search, allowing SNVs to be relocated and CNAs to be added or removed to maximize likelihood. This design naturally enforces longitudinal constraints while enabling modeling of sample-specific technical variation, such as differences in sequencing coverage and dropout rates across samples.

We apply LoPhy to longitudinal single-cell datasets comprising acute myeloid leukemia (AML) samples profiled with the Tapestri^®^ platform (Mission Bio, Inc.), including a cohort of 15 AMLs undergoing diverse treatments and four TP53-mutated AMLs sampled before and after therapy. LoPhy’s orthogonally-validated clone trees reveal a coherent picture of how these cancers evolved over time. These reconstructions reveal that AML clones emerging after therapy or at relapse are often defined by both SNVs and CNAs. While CNAs are well established drivers of oncogenesis in AML [19, 20], our results suggest that modeling them alongside SNVs is essential for fully characterizing key clonal populations.

## Materials and methods

LoPhy reconstructs clone trees from single-cell amplicon sequencing data collected at multiple time points from the same cancer. An overview of the method is provided in Figure 1. This section details the algorithm’s main components: required sequencing inputs, clone tree representation, evolutionary model, likelihood model, and optimization strategy. Central to LoPhy is a new factorized tree reconstruction objective, which extends concepts from our previous work [21].

**Fig 1.**
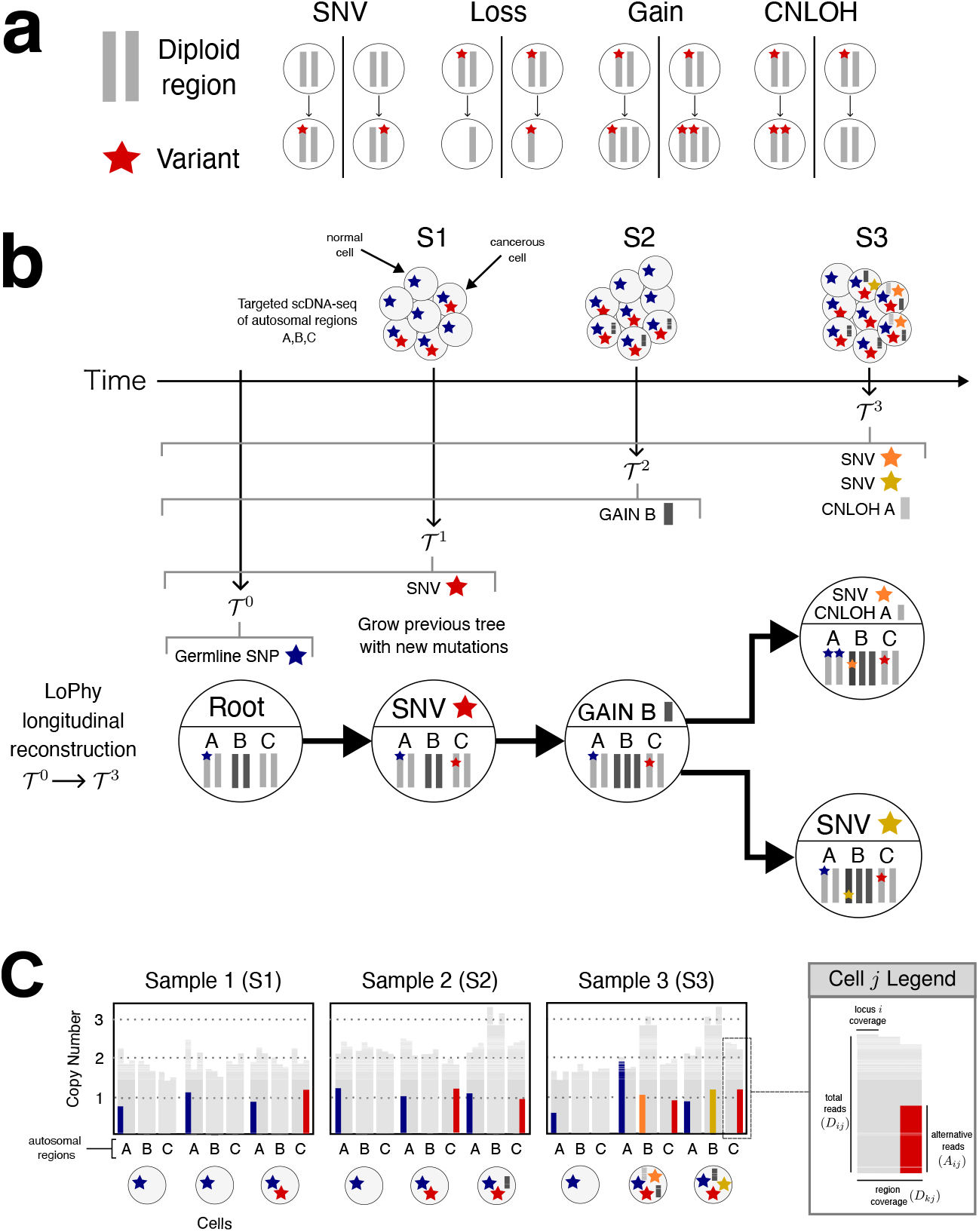
Overview of LoPhy. **a**. Four types of somatic events are modeled. **b**. Longitudinal reconstruction: LoPhy infers somatic events from longitudinal single-cell amplicon sequencing to build time-dependent subtrees 𝒯^1^, 𝒯 ^2^, 𝒯^3^, representing evolution up to time of sampling. **c**. Example for select cells from longitudinal sampling illustrating that single-cell amplicon sequencing provides region-level coverage and locus-level alternative/reference read counts.

### Inputs from longitudinal single-cell amplicon sequencing

Single-cell amplicon sequencing focuses on small genomic regions (150-450 base pairs) within disease-related genes. Amplicons—generated by amplifying the reverse transcription products of these regions—are used to measure the frequency of the target regions in the genome and identify variants. For each cell *j*, the data from a single-cell amplicon sequencing experiment is summarized as follows: *D*_*j*_ is the total number of reads mapped across all targeted regions, *D*_*kj*_ is the number of reads mapped to region *k, D*_*ij*_ is the total number of reads mapped to locus *i*, and *A*_*ij*_ is the number of alternative reads observed at locus *i*. Each cell is associated with a specific longitudinal sample, allowing the data to be partitioned into sample-specific subsets. We use **D** and **A** to denote the full dataset, containing read counts for all cells across all longitudinal samples. For a specific longitudinal sample *s*, we use **D**^*s*^ and **A**^*s*^ to represent the corresponding subset of data for cells originating from that sample.

### Clone tree representation

Inferring a clone tree that includes both SNVs and CNAs involves reconstructing a rooted, directed tree 𝒯, and clone assignment vector *σ*. Each node *v* in 𝒯 represents a distinct clone, with the root corresponding to the germline (non-cancerous) clone, which may contain germline SNPs but no somatic mutations. The clone assignment vector *σ* specifies the clone membership of each cell, such that if *σ*_*j*_ = *v*, cell *j* is assigned to clone *v*. The copy number profile of cell *j* is determined by its assigned clone *v*, where *c*_*k*_(*σ*_*j*_) denotes the total copy number for target region *k*, and 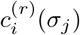 and 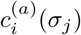 denote the number of copies of the reference and alternative alleles, respectively, at locus *i*.

### Evolutionary model

The LoPhy algorithm models four types of somatic events: SNV acquisition, copy number gains, copy number losses, and copy-neutral loss of heterozygosity (CNLOH). CNAs may affect either individual target regions or entire chromosomes. Each variant locus lies within a target region, and its copy number can change if a CNA impacts the allele containing that region. Because targeted sequencing data are noisy and offer limited resolution for detecting multiple copy number changes along the same lineage, we restrict each region to be impacted by at most one CNA per lineage in the tree. SNV evolution in 𝒯 follows the *k*-Dollo model [22], in which each SNV is acquired exactly once but may be lost at most once per lineage due to a deletion of the genomic region on which it resides.

### Longitudinal likelihood model

Given *S* longitudinal samples, LoPhy assumes the following factorized longitudinal likelihood:

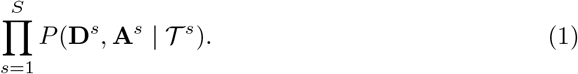

Here, each clone tree 𝒯^*s*^ represents the evolutionary history up to time point *s* and forms a subtree of all subsequent trees, 𝒯^1^*⊆* 𝒯^2^⊆· · · ⊆𝒯^*S*^. This factorization relies on two key assumptions: (1) somatic mutations accumulate over time, and (2) any mutation that arises between time points *s−*1 and *s* is carried by one or more cells sampled at time point *s*. Assumption (1) is strongly supported in many cancer types, where clonal evolution is largely tree-like [4, 23]. Assumption (2) is valid as the number of cells sampled at each time point increases. Under these assumptions, if a mutation is first detected at time point *s*, it must be present in at least one clone in 𝒯^*s*^ and absent from all clones in 𝒯^*s−*1^, which naturally leads to the nested tree structure.

The factorization in Eq. 1 approximates the joint likelihood over the data from all *S* longitudinal samples:

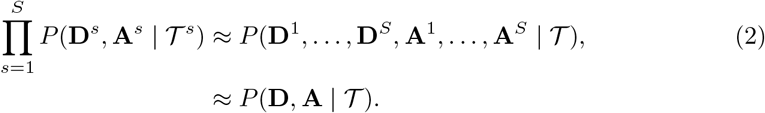

The left and right hand sides of Eq. 2 are equivalent if and only if

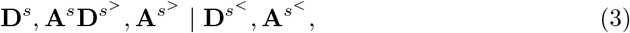

which states that the read count data for sample *s* are conditionally independent of all data from future samples, *s*^*>*^, given the data from all prior samples, *s*^*<*^. This conditional independence follows directly from assumption (2): if all mutations present up to time point *s* are captured through longitudinal sequencing, then the subtree 𝒯^*s*^ is fully determined by the data collected at or before *s*, making Eq. 1 a valid approximation of the joint likelihood.

Each factor in Eq. 1, which we refer to as the *sample-specific likelihood*, is defined as:

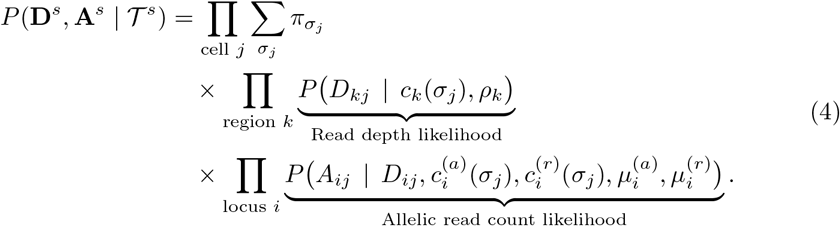

The sample-specific likelihood marginalizes over all possible clonal assignments for *σ*_*j*_ defined by 𝒯^*s*^, with 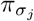 denoting the prior probability of assignment to clone *σ*_*j*_. The read depth likelihood, which evaluates the inferred total copy numbers for each target region in each cell, and the allelic read count likelihood, which evaluates the inferred number of alternative and reference allele copies at each locus in each cell, are described below.

### Read depth likelihood

Current single-cell amplicon sequencing technologies have non-uniform coverage across amplicons [12]. Since each amplicon covers a portion of a disease-specific gene (i.e., a region), the probability a read falls into region *k*, denoted with *ρ*_*k*_, is impacted by a variety of factors including the size of the region, its GC content, etc. We estimate *ρ*_*k*_ using the observed fraction of reads that fall into region *k* in a population of normal (i.e., absent of CNAs) cells. For a cell *j* with *D*_*j*_ total reads, assigned to clone *σ*_*j*_, and copy number *c*_*l*_(*σ*_*j*_) for region *l*, the expected read depth in region *k* is given by

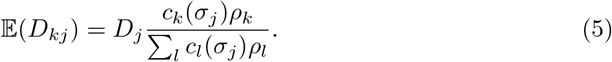

Cells assigned to the same clone may have larger than expected variances in their regional read depths, a phenomenon known as *overdispersion*. To model this, we use a Negative Binomial distribution for each clone’s regional read counts, parameterized by the mean *µ* = 𝔼 (*D*_*kj*_) and inverse-dispersion parameter *θ*:

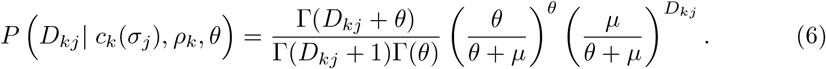

This parametrization, often referred to as the *overdispersed Poisson*, differs from the standard Poisson in that its variance is 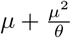, while the standard Poisson has variance equal to *µ*. Setting *θ* =*∞* assumes no overdispersion and reduces it to the Poisson, while smaller values of *θ* account for greater overdispersion. We omit *θ* from the read depth likelihood in Eq. 4, since it is treated as fixed.

### Allelic read count likelihood

Although single-cell amplicon sequencing has low coverage, allelic imbalances resulting from CNAs can still be detected from shifts in the observed proportions of alternative and reference read counts. However, inferring the relative proportions of alternative and reference alleles from these read counts is complicated by overdispersion, which arises from allelic biases during amplification. To address this, the allelic read counts are modeled with a beta-binomial distribution. Using this model, the likelihood of observing *A* alternative reads out of *D* total reads at a position, given *c*^(*a*)^ alternative alleles and *c*^(*r*)^ reference alleles, is:

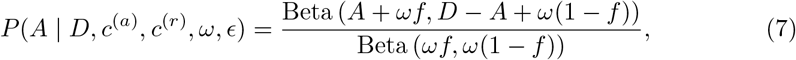

where *ω >* 0 is the beta-binomial concentration parameter, Beta(*·*) denotes the Beta function, and *f* is the expected fraction of alternative reads assuming a fixed sequencing error rate of *ϵ*:

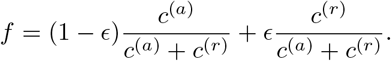

Allelic read counts are also impacted by allele-specific dropout during sequencing, leading to false negatives. To account for this, we introduce global allele-specific dropout rates 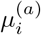 and 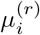, representing the dropout probabilities of the alternative and reference alleles at locus *i*. The number of amplified alleles can be modeled using a binomial distribution. Specifically, the likelihood of amplifying *l* alternative and *k* reference alleles at locus *i*, given 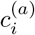 alternative and 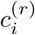 reference alleles copies, is:

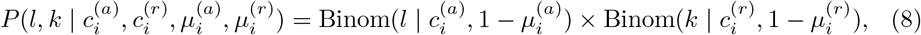

where Binom 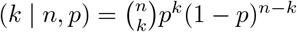 is the binomial likelihood of observing *k* successes out of *n* trials with a success rate of *p*. To fully account for dropout uncertainty, the likelihood of the observed read count *A* is obtained by summing over all possible dropout scenarios:

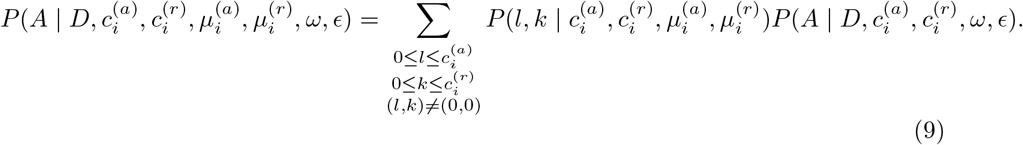

For succinctness, we omit the fixed parameters *ω, ϵ*, defining the shorthand used in Eq. 4:

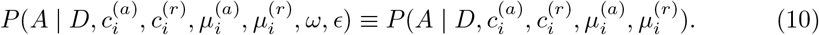

### Tree prior

Single-cell sequencing data are noisy, and complex evolutionary models can overfit the data by introducing spurious clones or CNAs. To mitigate this, we impose a prior on the number of clones and CNAs in the inferred tree.

Each CNA event incurs a penalty, with CNLOH assigned a stronger penalty than a gain or loss. CNAs affecting contiguous regions are counted as a single event, as large-scale alterations (e.g., whole-chromosome changes) are common. We also penalize the total number of clones, since incorporating more SNVs and CNAs into the tree increases the chance that multiple mutations originated in the same clone. Both penalties scale linearly with the number of cells, as larger datasets exert a proportionally greater influence on the likelihood. Let *p*_1_ denote the penalty for clones and *p*_2_ the penalty for CNAs. The prior is defined as:

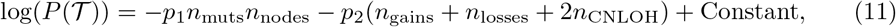

where *n*_muts_, *n*_nodes_, *n*_gains_, *n*_losses_, and *n*_CNLOH_ denote, respectively, the number of mutations, clones, non-contiguous gains, non-contiguous losses, and non-contiguous CNLOH events.

### Algorithm

The LoPhy algorithm combines sequential tree-building with stochastic search to generate a sequence of clone trees 𝒯 ^1^, …, 𝒯^*S*^ that maximize the factorized posterior:

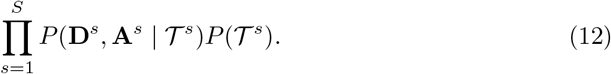

Each factor in the posterior is the product of the prior (Eq. 11) and sample-specific likelihood (Eq. 4). The algorithm consists of three stages.

### Hyperparameters

Normal (non-cancerous) cell populations are first identified in each sample to estimate region-specific coverage probabilities *ρ*_*k*_ for each region *k*. LoPhy does this by independently reconstructing an SNV-only tree for each longitudinal sample, maximizing the sample-specific likelihood in Eq. 4, and designating cells assigned to the root node as the normal population. Finally, the sample in which each SNV is first detected is determined from the single-cell sequencing inputs.

### Tree initialization

The tree 𝒯^0^ is initialized with a root node representing the normal cell population, which may have germline SNPs, but no somatic mutations.

### Stochastic search

For each sample *s*, SNVs first observed in that sample are incorporated into 𝒯^*s−*1^ by forming new clones that descend from existing ones. After introducing new SNVs, stochastic search proceeds for *η* epochs. At each iteration, either a newly added SNV or a genomic region is selected at random, and all valid modifications are evaluated. Three types of moves are permitted: (1) relocate an SNV, (2) remove a CNA, or (3) add a CNA. CNA moves are constrained so that only regions in lineages not previously altered by a CNA in 𝒯^*s−*1^ may be modified. All tree modifications respect longitudinal constraints: clones in 𝒯^*s−*1^ remain fixed, and new SNVs/CNAs are restricted to descendant clones. Tree modifications are accepted if they increase the posterior likelihood (Eq. 12).

### Inferring clone assignment priors and dropout rates

Dropout rates and clone assignment priors are estimated for each sample *s* using its corresponding tree 𝒯^*s*^. We adopt the expectation–maximization (EM) algorithm from [12] to jointly estimate the prior probability of assignment to each node (clone) *v, π*_*v*_, and the global dropout rates for the alternative and reference alleles at each locus *i*, 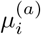 and 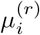. We place a Beta(*α, β*) prior on 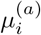 and 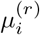 centered at 0.05, taking advantage of its conjugacy with the binomial likelihood used to model dropout rates.

For clone assignments, we use a flat Dirichlet(1, …, 1) prior, which is conjugate to the categorical distribution for clone assignment prior probabilities. The EM algorithm is run after every tree modification during stochastic search. There are two types of latent variables for each cell *j*: the clone assignment *σ*_*j*_, and the number of amplified alternative and reference alleles for each locus *i*, 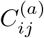 and 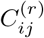. Given the current parameter estimates, the E-step is defined as:

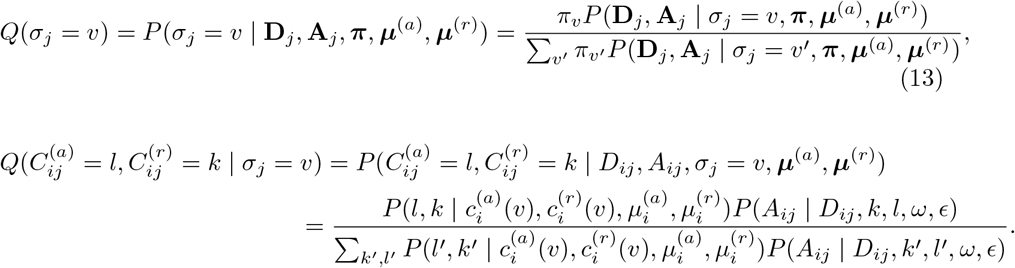

The update equations for the M-step are defined as:

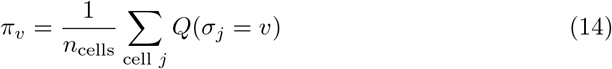

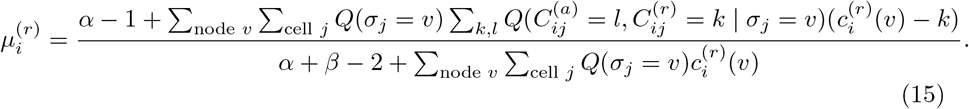

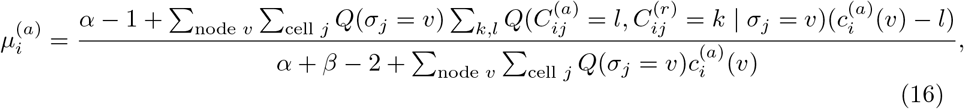

## Results

### Evaluation on simulated longitudinal data

We benchmarked LoPhy on simulated longitudinal cancer data, comparing it to COMPASS [12], SCITE [24], and LACE [14]. COMPASS reconstructs clone trees incorporating both SNVs and CNAs but does not account for longitudinal sampling. SCITE is a widely used method for SNV-based tree reconstruction, though it is not designed for longitudinal data. LACE is designed for longitudinal data but models only SNVs. We also considered BiTSC^2^ [11], which supports joint reconstruction of SNVs and CNAs, but excluded it due to prohibitively long runtimes, even on small longitudinal datasets. Because COMPASS and SCITE do not model longitudinal sampling, we provided them with pooled data consisting of cells from all time points, whereas LoPhy and LACE were given the time point at which each cell was sampled. We simulated longitudinal cancer datasets to assess how jointly modeling SNVs and CNAs in a longitudinal framework improves clone tree reconstruction, compared to methods that either model only SNVs or are not designed for longitudinal data. We simulated 40 longitudinally observed cancers in total: 10 for each setting of longitudinal samples (2, 3, 4, or 5). Each dataset spanned 2–5 time points, with 3,000 cells sampled per time point (6,000–15,000 cells per dataset). Sequencing data were generated to resemble output from the Tapestri ^®^ platform (see Section A3 in S1 Appendix), comprising 20 genomic regions, 20 SNVs, and 3 CNAs. Ground-truth clone trees included 6–9 clones, with the number of clones increasing in datasets with more time points, reflecting the expectation that more longitudinal sampling reveals greater clonal diversity. The longitudinal dynamics for each simulated cancer were generated using a Moran process [25], a well-established model in population genetics.

Performance was evaluated using cell-to-clone assignment accuracy—comparing the inferred mutant (alternative) and total copy number profiles of each cell to its ground-truth profile—and by measuring the correspondence between the inferred and ground-truth clone tree structures. Specifically, we computed the mean absolute error (MAE) between inferred and ground-truth mutant copy numbers (MCN) and between inferred and ground-truth total copy numbers (TCN), denoted MCN-MAE and TCN-MAE, respectively. We also introduce a new metric, the false emergence rate (FER), that measures the fraction of cells assigned copy numbers found exclusively in clones from future samples and absent from all clones detected up to the cell’s sampling time. FER captures a key pitfall of ignoring longitudinal structure: mutations may appear to emerge earlier in evolution than they actually did. Tree structure correspondence is measured by the Tree F1 score, which summarizes the precision and recall of evolutionary relationships between pairs of mutations in the inferred tree relative to those in the ground-truth tree. The metric evaluates SNV–SNV, CNA–CNA, and SNV–CNA mutation pairs. In addition to the Tree F1 score, in Section A4 in S1 Appendix we report precision, recall, and F1 scores for each mutation–pair category (SNV–SNV, CNA–CNA, SNV-CNA). Detailed descriptions of all evaluation metrics are provided in Section A2 in S1 Appendix.

Figure 2 summarizes the performance of LoPhy and competing methods on the 40 simulated longitudinal cancer datasets. LoPhy, on average, achieved the best performance across all evaluation metrics, regardless of the number of longitudinal samples, and showed consistently strong results—particularly for total copy number estimation and tree structure inference. In contrast, COMPASS struggled with total copy number inference and often introduced erroneous CNAs, leading to an average TCN-MAE above 0.35 across all simulation settings and poor correspondence with the ground-truth tree structures. This behavior is expected, as COMPASS does not model coverage variation across longitudinal samples, causing CNAs to be inferred from differences in coverage between samples rather than true copy number changes. LoPhy also achieved the best average performance for mutant copy number inference.

**Fig 2.**
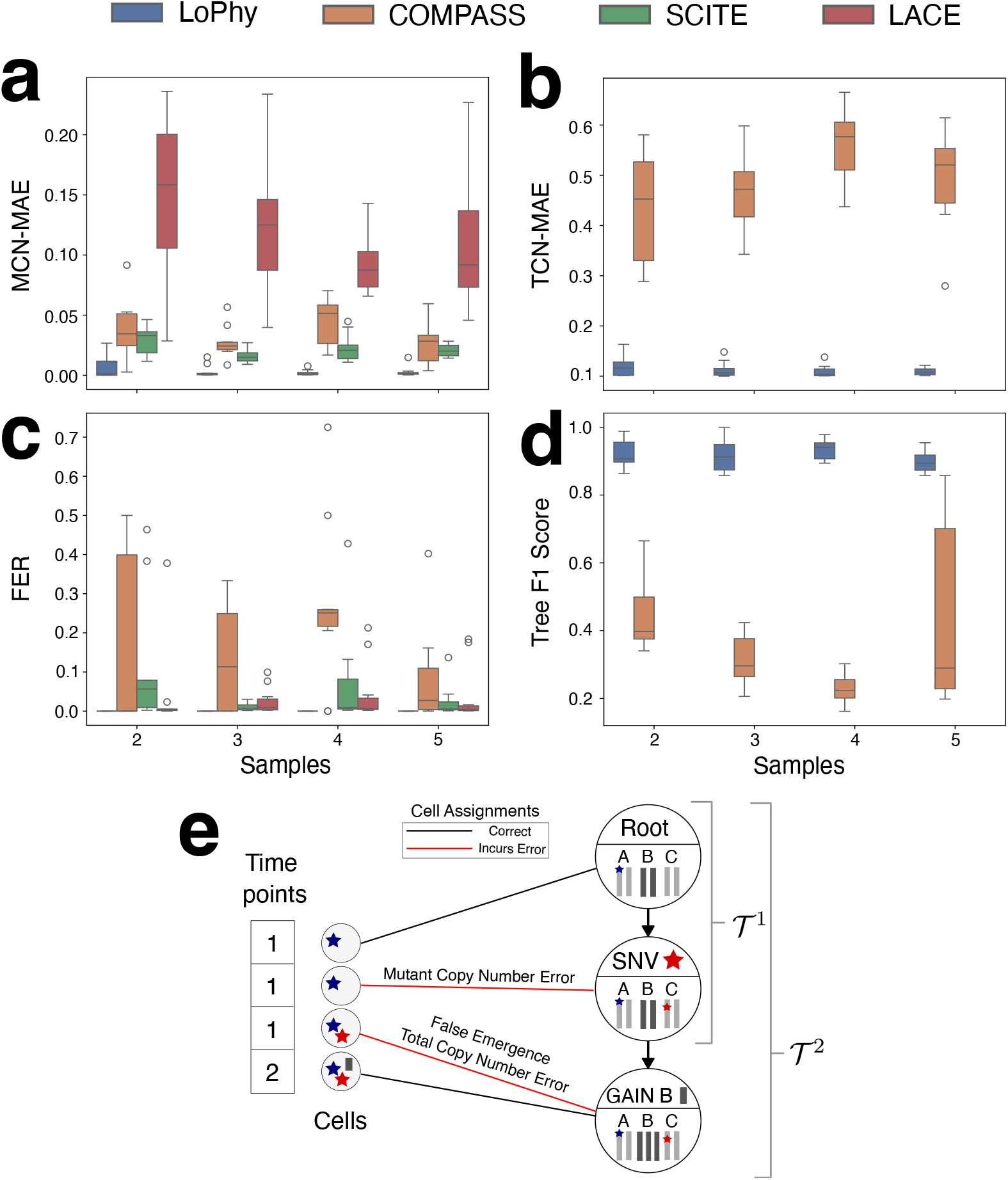
Evaluation of LoPhy, COMPASS, SCITE, and LACE on 40 simulated longitudinal cancer datasets. **a-d**. Distributions of evaluation metrics: MCN-MAE (*mutant copy number mean absolute error*), TCN-MAE (*total copy number mean absolute error*), FER (*false emergence rate*), and Tree F1 score. Box plots indicate the median (center line), first and third quartiles (box edges), range of data (whiskers), and outliers (unfilled circles). For MCN-MAE, TCN-MAE, and FER (a–c), lower values indicate more accurate tree reconstructions and cell assignments. For the Tree F1 score (d), higher values indicate closer agreement between the inferred and ground-truth tree structures. **e**. Examples of cell assignments illustrating the three error different types.

Although COMPASS and SCITE showed slightly lower MCN-MAE values on two datasets, the difference was very small—approximately 7 *×* 10^*−*4^.

SCITE estimates whether loci are heterozygous or homozygous but does not infer total copy numbers; therefore, we report MCN-MAE for SCITE but exclude it from TCN-MAE and Tree F1 score comparisons. LACE outputs a binary matrix indicating the presence or absence of SNVs per cell, so we include it in the MCN-MAE comparisons while noting this limitation; it is likewise excluded from TCN-MAE and Tree F1 score. SCITE performed well on MCN-MAE, showing lower error than COMPASS on average across all simulation settings. LACE exhibited poor MCN-MAE performance, likely due to its binary output, but performed well on the FER metric, as expected given its longitudinal modeling. LoPhy achieved perfect FER, whereas COMPASS frequently inferred mutant and/or total copy number changes earlier in evolution than they actually occurred.

Finally, LoPhy was substantially faster than COMPASS, completing datasets with 5 longitudinal samples in an average of about 9.5 minutes, compared to roughly 1 hour and 40 minutes for COMPASS. Runtime comparisons for all methods across the different simulation settings are provided in Section A4 in S1 Appendix.

### Longitudinal phylogenetic analysis of acute myleoid leukemias

We applied LoPhy to a cohort of 15 longitudinally observed acute myeloid leukemias (AMLs) profiled with the Tapestri^®^ platform [8], each with 2–5 time points sequenced using either a 50-amplicon panel (19 genes) or a 279-amplicon panel (37 genes). The authors in [8] provided clinical descriptions for all of these AMLs, which are summarized in Section A1 in S1 Appendix, detailing treatments and changes in the clonal composition of the AMLs over time. We demonstrate that our analyses with LoPhy are consistent with these clinical descriptions, and reveal that AML clones dominant after treatment or at relapse frequently harbor both SNVs and CNAs, illustrating the clinical relevance of jointly modeling SNVs and CNAs in a longitudinal framework. Below, we highlight several representative cases, including AML-99 and two AMLs where we directly compare reconstructions from LoPhy and COMPASS. Analyses of the remaining AMLs, including a second cohort of four TP53-mutated AMLs with samples collected before and after venetoclax treatment [10], are provided in Section A5 in S1 Appendix.

#### Dominant AML clones are characterized by both SNVs and CNAs

AML-99 illustrates how incorporating longitudinal information and jointly modeling SNVs and CNAs yields more biologically meaningful clone trees. This patient was diagnosed with AML carrying an IDH2 R140Q mutation (AML-99-001) and had two additional samples collected during therapy (AML-99-002, AML-99-003). The patient remained in remission until relapse (AML-99-004), after which therapy was adjusted, though the disease remained refractory (AML-99-005).

Figure 3a shows the longitudinally-observed clone tree reconstructed by LoPhy for AML-99, revealing that clones defined by CNAs dominated as the disease evolved. Across the first three samples (AML-99-001 through AML-99-003), the dominant clone carried a MYC/RAD21 gain and CNLOH of the RUNX1 reference allele, consistent with the bulk ASCAT [26] profile for AML-99-001 (Figure 3b) from [12]. At relapse, however, a new clone emerged with the same MYC/RAD21 gain but CNLOH of the RUNX1 alternative allele, dominating AML-99-004 (Figure 3c). After therapy was adjusted, the relapse clone contracted, while a FLT3-positive subclone of the original MYC/RAD21 clone expanded to dominate AML-99-005 (Figure 3c).

**Fig 3.**
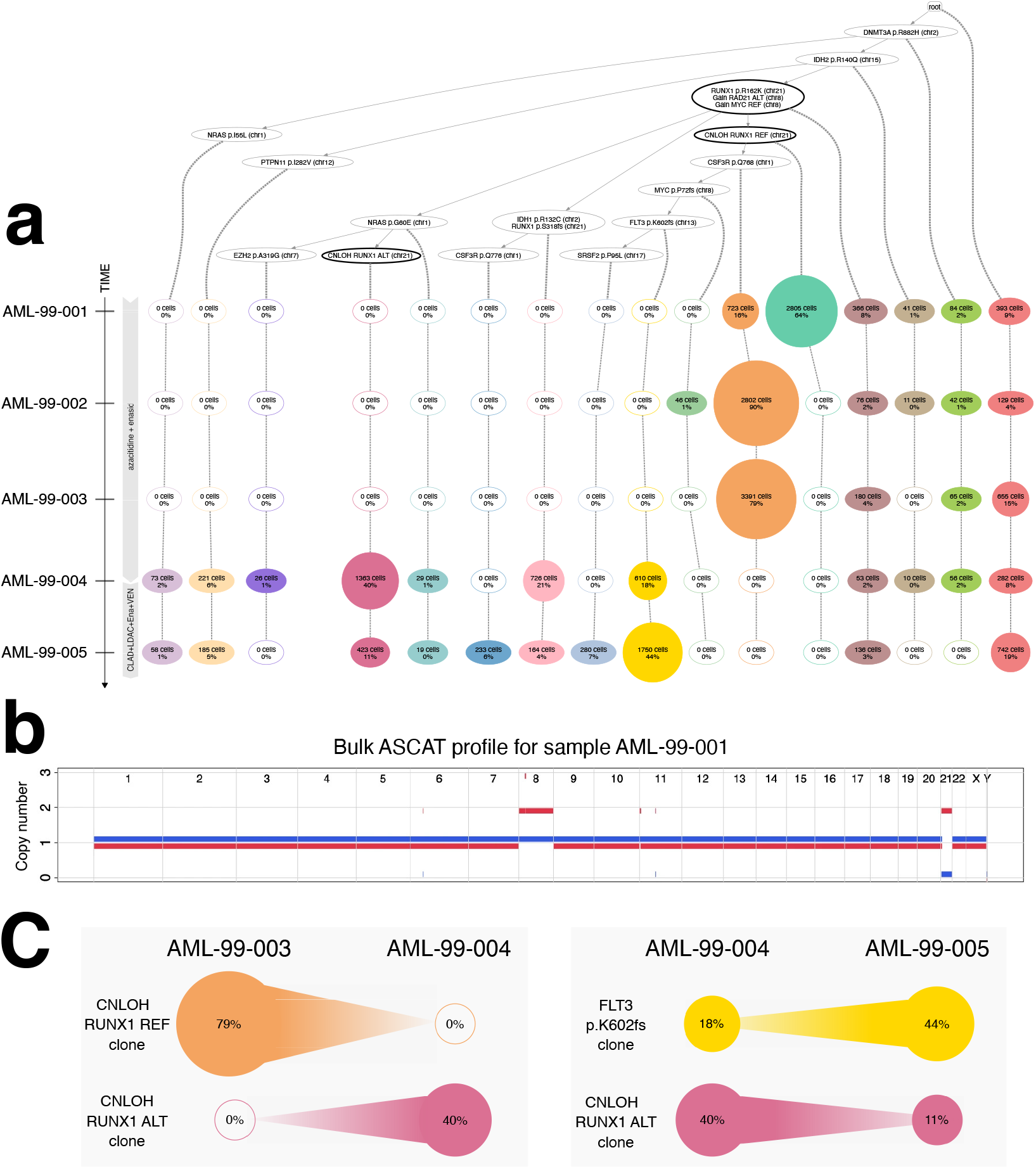
**a**. Longitudinally-observed clone tree reconstructed by LoPhy for AML-99. Nodes containing CNAs are shown with bold outline. The time axis indicates sampling points and therapy regimens: Clad, cladribine; LDAC, low-dose cytarabine; Ena, enasidenib; VEN, venetoclax; DAC, decitabine. **b**. ASCAT profile for sample AML-99-001 from [12]. **c**. Fluctuations in size of key clonal populations in response to therapy.

A key insight from this reconstruction is that two independent CNLOH events occurred in the RUNX1 region on separate lineages in the tree: one deleting the reference allele early in the disease, and one deleting the alternative allele that appears only at relapse. At relapse, the lineage with CNLOH of the RUNX1 alternative allele expanded and became dominant, while the earlier lineage carrying CNLOH of the reference allele contracted (Figure 3c). The CNLOH at relapse is well supported by the data—cells in this lineage have median coverage *≥* 30 at the RUNX1 p.R162K locus, yet nearly all have zero (or near-zero) reads supporting the alternative allele. Bulk data for AML-99-004, if available, could independently confirm the CNLOH event at relapse.

We provide COMPASS’s reconstruction for AML-99 in Section A5 in S1 Appendix, which agrees poorly with the clinical description for this patient and infers CNAs which cannot be validated. Across both cohorts—the 15 AMLs from [8] and the 4 TP53-mutated AMLs from [10]—LoPhy reconstructed clone trees where CNA-defined clones dominated in 17 of 19 AMLs. Bulk data exists for 5/17 of these CNA-dominated AMLs, supporting the majority of CNAs inferred by LoPhy.

#### Comparing and contrasting LoPhy and COMPASS longitudinal AML reconstructions

Figure 4 shows longitudinal reconstructions by LoPhy and COMPASS for AML-01 and AML-83. These two cases highlight the importance of longitudinal constraints and estimating sample-specific coverage and dropout rates.

**Fig 4.**
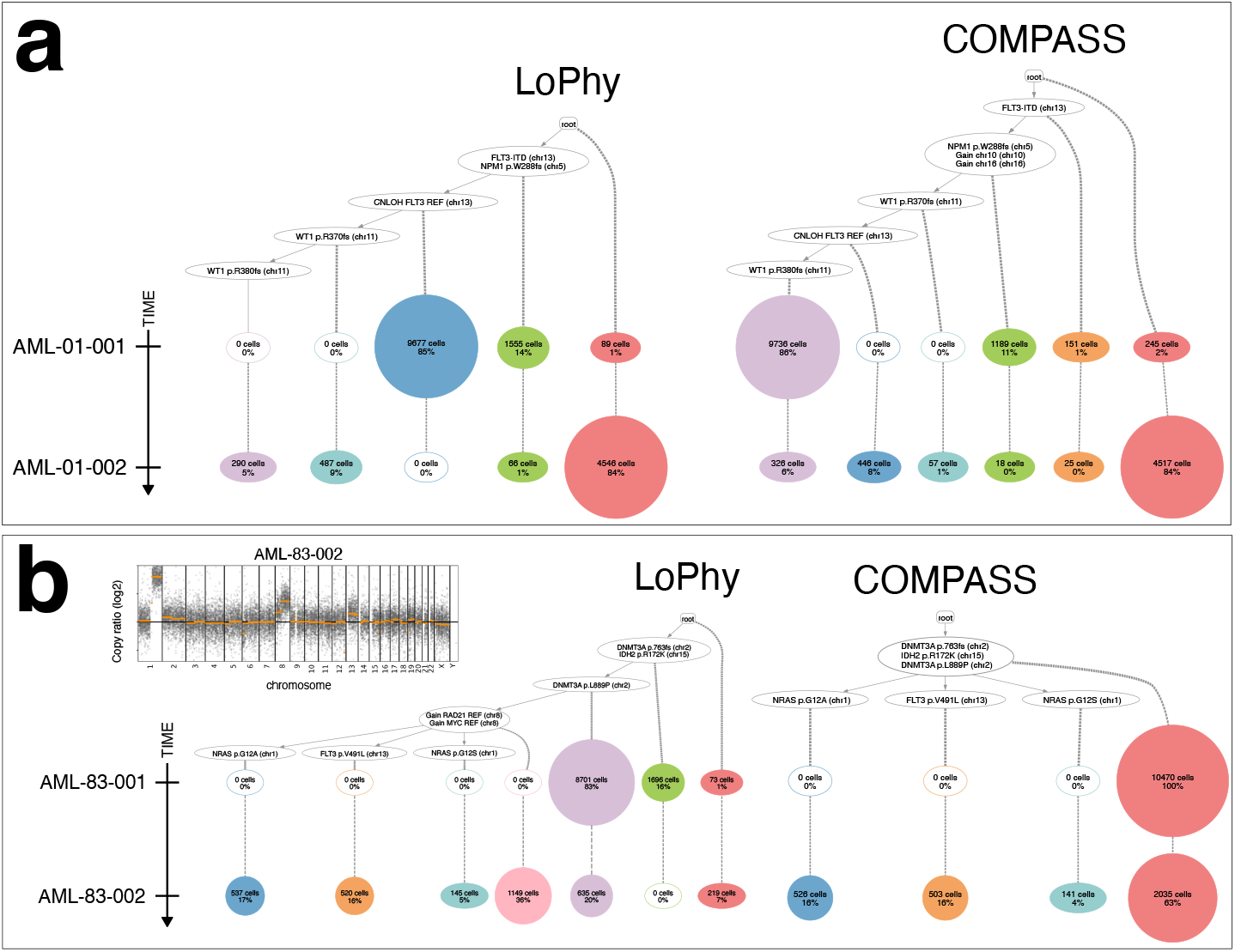
Longitudinal reconstructions by LoPhy and COMPASS for AML-01 (**a**), and AML-83 (**b**).

Figure 4a shows the longitudinally-observed clone trees reconstructed by LoPhy and COMPASS for AML-01. This is a FLT3-ITD mutated AML with samples collected at diagnosis (AML-01-001) and relapse (AML-01-002). Treatment was given after diagnosis; however, the relapse sample remained positive for FLT3-ITD [8]. Although both LoPhy and COMPASS infer a CNLOH of the FLT3 reference allele, their reconstructions differ in its placement. LoPhy places the FLT3 CNLOH before the two WT1 mutations, identifying 85% of cells at diagnosis as positive for this event. COMPASS places the CNLOH event after the first WT1 mutation, assigning 86% of cells at diagnosis both WT1 mutations and the CNLOH event—even though no WT1 mutations were detected at diagnosis. This high rate of false emergences occurs because, without WT1 mutation data at diagnosis or longitudinal constraints, COMPASS relies solely on the strong CNLOH signal when assigning cells to clones. This also results in COMPASS estimating unrealistically high dropout rates of 50% for the WT1 loci, highlighting the importance of sample-specific dropout modeling for plausible longitudinal reconstructions.

Figure 4b shows the trees reconstructed by LoPhy and COMPASS for AML-83. This AML is characterized at diagnosis (AML-83-001) by an IDH2 p.R172K mutation. At relapse (AML-83-002), two additional NRAS mutations were detected that were absent at diagnosis [8]. Bulk copy number analysis using CNVkit [27], as reported in [12], supports a copy number gain on chromosome 8 at relapse (Figure 4b top left). LoPhy reconstructs an evolutionary history consistent with the case description in [8], correctly identifying the IDH2 mutation at diagnosis, and the NRAS mutations and MYC/RAD21 gain on chromosome 8 at relapse. The MYC/RAD21 clone dominates at relapse, giving rise to three subclones distinguished by distinct FLT3/NRAS SNVs. In contrast, while COMPASS generally recovers the correct SNV evolution, it misses the MYC/RAD21 gain. This occurs because it does not model sample-specific coverage, so differences between the two sequencing experiments—particularly among the “normal” cells used to define baseline coverage—distort the signal needed to detect the gain.

When run only on the relapse sample (AML-83-002), where sample-specific coverage does not need to be modeled, COMPASS reconstructs a tree nearly identical to LoPhy’s, including the MYC/RAD21 gain. However, analyzing the two AML-83 samples independently with COMPASS produces inconsistent clone structures across the trees, obscuring the temporal ordering of genomic changes and the relationships between cells from the two samples.

Analyses of both AML cohorts with LoPhy—including the 15-AMLs from [8] and the 4 TP53-mutated AMLs from [10]—provide the first integrated view of how SNVs and CNAs jointly shaped clonal evolution in these AMLs over time. These remaining analyses, presented in Section A5 in S1 Appendix, mirror the findings for AML-01 and AML-83. COMPASS inferred trees that were inconsistent with longitudinal data or clinical descriptions in 7 of 19 AMLs, whereas LoPhy produced consistent reconstructions in all but one case, highlighting the robustness of its modeling framework. Note that previous analyses of these AMLs were limited either to SNV-only reconstructions or to reconstructing trees independently for each time point. In the original study [8], the 15 longitudinal AMLs were analyzed with SCITE [24] by pooling all time points and inferring clone trees based solely on SNVs; only AML-99 was later reanalyzed with COMPASS [12], which was used to reconstruct trees independently for each time point. The 4 TP53-mutated AMLs from [10] were likewise analyzed with COMPASS in [12], by reconstructing trees independently for each time point.

## Discussion

Longitudinal single-cell sequencing is increasingly used to study cancer evolution, particularly in leukemias and lymphomas. Methods exist to reconstruct clone trees from longitudinal single-cell data; however, they typically model the evolution of either single-nucleotide variants (SNVs) or copy number alterations (CNAs), but not both.

New amplicon-based single-cell DNA sequencing technologies now enable reliable detection of both SNVs and CNAs, allowing detailed longitudinal tracking of these genomic alterations at single-cell resolution. Here, we show that reconstructing clone trees from longitudinal single-cell amplicon sequencing data reveals that dominant leukemia clones are often defined by both SNVs and CNAs, and that these clones frequently expand or contract in response to therapy. These insights are made possible by our algorithm, LoPhy, which optimizes a novel factorized objective to reconstruct clone trees capturing the joint evolution of SNVs and CNAs from longitudinal single-cell data. We show that LoPhy reconstructions are consistent with clinical observations and supported by orthogonal bulk sequencing data from the same cancers. The resulting trees provide continuous joint histories of how SNVs and CNAs emerge and evolve over time within individual cancers. The accuracy of our algorithm is further supported by simulations, where LoPhy outperformed existing reconstruction methods across multiple evaluation metrics. We further demonstrate in Section A4 in S1 Appendix that LoPhy accurately reconstructs clone trees from single–time point data, showing that it is applicable to both single–time point and longitudinal settings.

The current LoPhy algorithm has a few notable limitations. First, its longitudinal likelihood model assumes that sequencing adequately captures mutations present at each time point, which may not hold if the sequencing quality is low or if the sampled cells do not fully represent the true clonal populations. While this has not posed issues in the available single-cell amplicon sequencing datasets, additional modeling of cross-sample dropout could be needed to account for missing mutations. Second, LoPhy uses a greedy search strategy, which can become trapped in local maxima. Although targeted single-cell experiments typically profile thousands of cells, sparsely sampled subclones could be missed under this approach. This issue may be mitigated by increasing the number of cells sampled at each time point. Lastly, our copy number evolution model is potentially too simplistic: it only allows each region to be impacted by at most one CNA per lineage, models a restricted set of CNA types, and does not account for homozygous loss—a rare but possible event.

Future work could address these limitations. A more comprehensive copy number model that incorporates homozygous loss and multiple CNAs affecting the same region within a lineage is feasible, though these additions may become more relevant as single-cell technologies advance to better capture such events. The search strategy could be improved by replacing the greedy approach with a Markov Chain Monte Carlo framework, reducing the risk of becoming trapped in local maxima. However, this would require a carefully designed proposal distribution and acceptance criteria to avoid introducing spurious CNAs. Finally, while LoPhy was developed for single-cell amplicon sequencing, the framework could be extended to reconstruct longitudinally-observed clone trees from single-cell whole-genome sequencing or bulk sequencing.

## Supporting information

**S1 Appendix. Supplementary Information**

## Acknowledgments

Q.M. was partially supported by a NIH/NCI Cancer Center Support Grant P30 CA008748 (Vickers). The funders had no role in study design, data collection and analysis, decision to publish, or preparation of the manuscript.

