## Supplement for "Longitudinal Phylogenetic Inference of Copy Number Alterations and Single Nucleotide Variants from Single-Cell Sequencing"

#### A1 Detailed overview of longitudinal acute myeloid leukemia (AML) data

We applied LoPhy to two longitudinal cohorts: 15 AMLs with 2–5 samples collected during various treatments [1], and 4 TP53-mutated AMLs sampled before and after venetoclax treatment [2]. Below, we briefly summarize the studies these AMLs originated from and provide tables describing each longitudinal sample.

The authors in [1] collected between 1-5 samples from 123 AMLs to characterize the genotypes and phenotypes of AML. Of these 123 patients, 15 had longitudinal samples. These samples were profiled with the Tapestri® platform [1], using either a 50-amplicon (19-gene) or 279-amplicon (37-gene) panel. Table A1 provides annotations for each sample from these 15 longitudinally observed AMLs. The annotations are derived from the case descriptions provided in the supplemental materials of [1].

**Table A1.** Description of longitudinal samples for 15 AMLs from [1].

| ID | 01 | 02 | 03 | 04 | 05 |
|---|---|---|---|---|---|
| AML-01 | **Diagnosed** with therapy related FLT3-ITD AML with minimal maturation | **Relapsed** after therapy (cladribine, cytarabine, decitabine) positive for FLT3-ITD | | | |
| AML-04 | **Refractory** FLT3-ITD therapy related-AML after therapy (decitabine, ruxolitinib) | After therapy (azacitidine, quizartinib) | **Refractory** after therapy (crenolanib) | | |
| AML-07 | **Diagnosed** with pure erythroid leukemia | **Relapsed** after therapy (cytarabine) | | | |
| AML-09 | **Diagnosed** with FLT3-ITD therapy related-AML | **Relapsed** despite complete remission after therapy (azacitidine, sorafenib) | | | |
| AML-18 | **Diagnosed** with AML with maturation | **Relapsed** 1 yr after therapy (clofarabine + idarubicin + cytarabine) | | | |

| | | | | | |
|---|---|---|---|---|---|
| AML-21 | **Diagnosed** with AML | **Relapsed** after therapy (clofarabine + idarubicin + cytarabine) | | | |
| AML-38 | Had therapy at prior institution (azacitidine, decitabine), then had recurrent AML after 1 cycle of therapy (cytarabine, quizartinib) | After therapy (cytarabine, quizartinib) | **Refractory** after therapy (cytarabine, quizartinib) | | |
| AML-39 | **Diagnosed** with myelodysplastic syndrome progressed into secondary AML | **Relapsed** after therapy (decitabine, was refractory to clorofarabine + idarubicin + cytarabine) | | | |
| AML-63 | **Diagnosed** with FLT3-ITD AML with minimal maturation | After therapy (decitabine, vosaroxin) | **Complete remission** after 1 cycle | **Relapsed** after additional cycle | **Complete remission** after continued therapy (decitabine, vosaroxin) |
| AML-66 | **Diagnosed** with AML | **Relapsed** after therapy (guadecitabine) | | | |
| AML-83 | **Diagnosed** with IDH2 p.R172k mutated AML | **Relapsed** after several different therapy regimens (azacitidine + enasidenib, decitabine + venetoclax) | | | |
| AML-88 | **Diagnosed** with FLT3-ITD and FLT3 p.D835V AML | **Complete remission** after therapy (3 cycles of: cladribine, idarubicin, cytarbine (CLIA) plus midostaurin) | **Complete remission** after allogeneic stem cell transplant | **Relapsed** 2 months after transplant positive for FLT3-ITD | **Complete remission** after therapy (decitabine, venetoclax, gilteritinib), was refractory to decitabine + quizartinib. |
| AML-99 | **Diagnosed** with IDH2 R140Q AML with maturation | After 2 cycles of therapy (azacitidine, enasidenib) | After 6 cycles of therapy (azacitidine, enasidenib) | **Relapsed** after 1 yr of remission | **Refractory** after salvage chemotherapy (cladribine, cytarabine, enasidenib, venetoclax) |
| AML-107 | Therapy related-AML refractory to cladribine, cytarabine + filgrastim, decitabine, and gemtuzumab ozogamicin | **Refractory** to therapy (3 cycles of decitabine + venetoclax) | | | |

In [2], the authors investigated the effectiveness of venetoclax in AMLs with TP53 abnormalities. Bone marrow biopsies were collected before and after a 7-day venetoclax treatment, and cells were profiled using the Tapestri® platform (Mission Bio, Inc.). Four TP53-mutated AMLs were non-responsive, with TP53-mutant clones expanding after treatment regardless of dosage. Samples from two non-responsive AMLs (CAL-012, CAL-030) were profiled with a 20-gene AML panel, and two

(CVC-001, CAU-001) with a 45-gene panel. Table A2 summarizes the samples from these four non-responsive cases.

**Table A2.** Description of longitudinal samples for 4 TP53-mutated AMLs from [2].

| ID | 01 | 02 |
|---|---|---|
| CAL-012 | **Pre-venetoclax** approximately 50% of sampled cancer cells were TP53 mutant with PTPN1 and KRAS | **Post-venetoclax** TP53 mutant clone expanded to about 80% of sampled cells |
| CAL-030 | **Pre-venetoclax** approximately 40% of sampled cancer cells were TP53 mutant with PTPN11 and N/KRAS | **Post-venetoclax** TP53 mutant clone expanded to about 60% of sampled cells |
| CVC-001 | **Pre-venetoclax** approximately 20% of sampled cancer cells were TP53 mutant | **Post-venetoclax** TP53 mutant clone expanded to about 22% of sampled cells |
| CAU-001 | **Pre-venetoclax** approximately 70% of sampled cancer cells were TP53 mutant with CALR | **Post-venetoclax** TP53 mutant clone expanded to about 80% of sampled cells |

#### A2 Evaluation metrics

We evaluate LoPhy and competing algorithms on simulated data using four metrics. The first two are the *mutant copy number mean absolute error* (MCN-MAE) and the *total copy number mean absolute error* (TCN-MAE). These two metrics measure the average difference between the inferred and ground truth copy number profiles per cell. Given an inferred tree $\mathcal{T}$ with corresponding cell assignments $\sigma$, and a ground truth tree $\hat{\mathcal{T}}$ with ground truth cell assignments $\hat{\sigma}$, the MCN-MAE and TCN-MAE are defined as:

$$\text{MCN-MAE}(\mathcal{T}, \sigma, \hat{\mathcal{T}}, \hat{\sigma}) = \text{MAE}(c^{(a)}, \sigma, \hat{c}^{(a)}, \hat{\sigma}) \tag{1}$$

$$\text{TCN-MAE}(\mathcal{T}, \sigma, \hat{\mathcal{T}}, \hat{\sigma}) = \text{MAE}(c^{(t)}, \sigma, \hat{c}^{(t)}, \hat{\sigma}) \tag{2}$$

where:

- $c^{(a)}$ and $\hat{c}^{(a)}$ are the inferred and ground truth alternative (mutant) allele copy numbers,
- $c^{(t)}$ and $\hat{c}^{(t)}$ are the inferred and ground truth total copy numbers.

Each $\text{MAE}(c, \sigma, \hat{c}, \hat{\sigma})$ term calculates the mean absolute error between the inferred and ground truth copy numbers:

$$\text{MAE}(c, \sigma, \hat{c}, \hat{\sigma}) = \frac{1}{n_{\text{cells}} \times d} \sum_{\text{cell } j} \sum_{\ell} |c_\ell(\sigma_j) - \hat{c}_\ell(\hat{\sigma}_j)| \,. \tag{3}$$

Here, $n_{\text{cells}}$ is the total number of cells, $d$ is the number of loci or regions (depending on the context), and $\ell$ indexes them.

The third evaluation metric, *false emergence rate* (FER), quantifies the extent to which an inferred longitudinal tree $\mathcal{T}$ and cell assignments $\sigma$ imply that clones emerged earlier in evolution than they actually did. FER measures the fraction of cells incorrectly assigned to clones that harbor mutations—alternative or total copy number changes—not observed in the cell's originating sample but observed only in future samples. Let cell $j$ be drawn from sample $s$, and let $\sigma_j$ be its assigned clone. Then, cell $j$ is considered a false emergence if, for any region k or locus i:

$$\mathbf{1}_{\mathrm{FE}}(j) = \begin{cases} 1 & \begin{array}{l} \exists k \text{ such that } c_k^{(t)}(\sigma_j) \notin \hat{c}_k^{(t)}(s) \text{ and } \exists s' > s \text{ where } c_k^{(t)}(\sigma_j) \in \hat{c}_k^{(t)}(s') \\ \text{or } \exists i \text{ such that } c_i^{(a)}(\sigma_j) \notin \hat{c}_i^{(a)}(s) \text{ and } \exists s' > s \text{ where } c_i^{(a)}(\sigma_j) \in \hat{c}_i^{(a)}(s') \end{array} \\ 0 & \text{otherwise.} \end{cases}$$

Here, $\hat{c}_k^{(t)}(s)$ and $\hat{c}_i^{(a)}(s)$ are the ground truth sets of unique total and alternative allele copy numbers observed in sample $s$ at region $k$ and locus $i$, respectively. These sets are constructed from the unique copy numbers assigned to cells in sample $s$ using the ground truth assignments $\hat{\sigma}$. The FER is then defined as:

$$\mathrm{FER}(\mathcal{T}, \sigma, \hat{\mathcal{T}}, \hat{\sigma}) = \frac{1}{n_{\text{cells}}} \sum_{\text{cell } j} \mathbf{1}_{\mathrm{FE}}(j). \quad (4)$$

The last metric, *Tree F1 score*, evaluates the correspondence between the inferred and ground truth tree structures based on the evolutionary relationships between mutation pairs. The Tree F1 score is defined as:

$$\mathrm{F1}(\mathcal{T}, \hat{\mathcal{T}}) = 2 \times \frac{\mathrm{Precision}(\mathcal{T}, \hat{\mathcal{T}}) \times \mathrm{Recall}(\mathcal{T}, \hat{\mathcal{T}})}{\mathrm{Precision}(\mathcal{T}, \hat{\mathcal{T}}) + \mathrm{Recall}(\mathcal{T}, \hat{\mathcal{T}})}. \quad (5)$$

The Precision$(\cdot)$ and Recall$(\cdot)$ terms are defined as:

$$\begin{aligned} \mathrm{Precision}(\mathcal{T}, \hat{\mathcal{T}}) &= \frac{|R(\mathcal{T}) \cap R(\hat{\mathcal{T}})|}{|R(\mathcal{T})|} \\ \mathrm{Recall}(\mathcal{T}, \hat{\mathcal{T}}) &= \frac{|R(\mathcal{T}) \cap R(\hat{\mathcal{T}})|}{|R(\hat{\mathcal{T}})|}, \end{aligned}$$

where $R(\mathcal{T})$ is the set of pairwise evolutionary relationships between mutations encoded by the tree. In our definition, $R(\mathcal{T})$ includes SNV-SNV, CNA-CNA, and SNV-CNA pairs. Each ordered mutation pair $(m_1, m_2)$ can have one of the following relationships:

1. *ancestor-descendant*: $m_1$ occurs in a clone that is ancestral to the clone containing $m_2$.
2. *co-clustered*: $m_1$ and $m_2$ occur in the same clone.
3. *separate lineages*: $m_1$ and $m_2$ occur in separate clones on distinct lineages of the tree.

### A3 Simulation framework

We simulate longitudinally-observed cancer data using an adaptation of the framework described in [3]. Our framework introduces several key innovations to more realistically capture longitudinal cancer evolution and the variability observed in real single-cell amplicon data generated by platforms such as Tapestri® (Mission Bio, Inc.).

First, we model cancer evolution as a stochastic process using a Moran process [4], a population genetics model that incorporates birth, death, and selection dynamics. Discrete time points are sampled along this evolutionary trajectory, and the set of detectable clones at each time point is represented by a subtree $\mathcal{T}^1, \ldots, T^{n_{\text{samples}}-1}$ of the full tree $\mathcal{T}$.

Second, we introduce additional parameters to increase inter-sample heterogeneity. Because real single-cell amplicon data show substantial variation in dropout rates and region-level coverage across samples, we simulate them independently for each sample.

Our simulation framework expects the following inputs:

- $n_{\text{samples}}$: the number of longitudinal samples in the simulation.
- $n_{\text{cells}}$: the number of cells in each longitudinal sample.

- $n_{\text{nodes}}$: the number of nodes (clones) in the full tree $\mathcal{T}$.
- $n_{\text{regions}}$: the number of regions in the simulation.
- $n_{\text{SNVs}}$: the number of single-nucleotide variants (SNVs) in the simulation.
- $n_{\text{CNAs}}$: the number of copy number alterations (CNAs) in the simulation.
- $n_{\Delta\text{nodes}}$: the minimum number of new nodes (clones) in each longitudinal sample.

Given these inputs, longitudinal cancer data are simulated using the following process:

1. Generate a random tree $\mathcal{T}$ with $n_{\text{nodes}}$ nodes by uniformly sampling a Prufer sequence from $[0, n_{\text{nodes}} - 1]^{n_{\text{nodes}}-2}$.
2. Assign each of the $n_{\text{SNVs}}$ SNVs independently to one of the $n_{\text{regions}}$ regions, each with probability $1/n_{\text{regions}}$.
3. Assign SNVs and CNAs to the non-root nodes, with each mutation independently assigned to a node with probability $1/(n_{\text{nodes}} - 1)$, and ensuring that each non-root node receives at least one mutation.
4. Starting from the root node, traverse $\mathcal{T}$ in a randomized manner (mixing depth-first and breadth-first strategies) to generate $n_{\text{samples}} - 1$ subtrees, $\mathcal{T}^1, \ldots, \mathcal{T}^{n_{\text{samples}}-1}$. For the first $n_{\text{samples}} - 1$ longitudinal samples, each subtree defines the set of clones used to simulate sequencing data. By construction, each subtree must also include at least $n_{\Delta\text{nodes}}$ more nodes than the previous subtree. The final sample contains all clones defined by $\mathcal{T}$.
5. **Repeat** steps 6-9 for each longitudinal sample $s = 1, \ldots, n_{\text{samples}}$:
6. Sample dropout rates for each variant from a Beta$(1, 19)$ distribution, then clip them to the range $[0.1\%, 20\%]$. This distribution has a mean of 5.0% and standard deviation of 4.75%, producing dropout rates consistent with those observed in real data [1, 3].
7. Sample region weights $\rho_k$ from Dirichlet$(\alpha, \ldots, \alpha)$, with $\alpha = 1.5$. The minimum weight is set to $1/(n_{\text{regions}} \times 5)$ to ensure sufficient coverage.
8. Simulate node (clone) probabilities $(\pi_1, \ldots, \pi_{n_{\text{nodes}}})$ for each longitudinal sample according to the Moran process described in the next paragraph.
9. For each of the $n_{\text{cells}}$ cells in sample $s$:
   (a) Sample which node the cell is attached to from Categorical$(\pi_1, \ldots, \pi_{n_{\text{nodes}}})$.
   (b) For each variant $i$:
      - Sample the number of reads covering locus $i$ from a Poisson distribution with rate parameter $\lambda = 20 \times \rho_i \times n_{\text{regions}}$, where 20 is the average sequencing depth.
      - Each allele copy is dropped out according to the dropout probability for locus $i$ from step 6. The mutant read counts are drawn from a Binomial distribution with total reads drawn from the Poisson distribution described in previous step, and the probability of a mutated read being drawn is set to $f = \frac{c^{(a)}}{c^{(r)}+c^{(a)}}(1-\epsilon) + \frac{c^{(r)}}{c^{(r)}+c^{(a)}}\epsilon$, where $c^{(a)}$ and $c^{(r)}$ are the number of amplified alternative and reference alleles at locus $i$, and $\epsilon$ is the sequencing error rate which we set to 0.02.

We simulate the node (clone) assignment probabilities for cancer cells in each sample using a *Moran process*—a stochastic model used to describe the dynamics of finite populations. This model has been shown to produce realistic simulations of cancer evolution under selection and therapeutic pressure [4]. In the Moran process, a population of $N$ cells undergoes repeated rounds of cell birth, death, and selection, leading to gradual shifts in clonal composition over time. At discrete time points, we record the clonal composition of the population and use it to derive node assignment probabilities. Specifically, we define $\pi_1$ as the probability of assigning a cell to be normal (non-cancerous), while $\pi_2, \ldots, \pi_{n_{\text{nodes}}}$ are the probabilities of being assigned to different cancer clones. The set of detectable cancer clones in each sample $s$ is assumed to known, corresponding to the nodes in the subtree $\mathcal{T}^s$ generated in step 4 of our simulation framework.

The evolution of the cancer cell population proceeds according to the Moran process as follows:

1. For each time point $s = 1, \ldots, n_{\text{samples}}$:
   (a) Initialize or update the population
      - If it's the first time point, then randomly assign the $N$ cells to cancer clones defined by $\mathcal{T}^1$.
      - If it's some time point $s > 1$, then introduce new cancer clone populations in $\mathcal{T}^s$ by randomly selecting a portion of the $N$ cells to assign to each new clone. We assigned 1% of the N cells to each new clone.

(b) For each new clone $c$, draw an initial fitness coefficient $w_c \sim \text{LogNormal}(\mu, \sigma)$. We set $\mu = \max(w_1, \ldots, w_{n_{\text{nodes}}})$ which ensures new clones are at least as fit as existing ones, and set $\sigma = 0.3$. Fitness coefficients are clipped in the interval $[0.1, 10]$.

(c) Evolve the population over a fixed number of epochs:

i. Update each clone's fitness coefficient by random drift: $w_c = w_c * \Delta$, where $\Delta \sim \text{LogNormal}(0, 0.01)$.

ii. Compute the probability of being assigned to each clone as $p_c = \frac{f_c w_c}{\sum_{c'=2}^{n_{\text{nodes}}} f_{c'} w_{c'}}$, where $f_c$ is the current frequency of clone $c$.

iii. With uniform probability $1/N$, select one cell from the current population to be removed (i.e., to die).

iv. Replace it with a new cell whose clone assignment is drawn from $\text{Categorical}(p_2, \ldots, p_{n_{\text{nodes}}})$.

(d) Record the final clone frequencies at time point $s$: $(f_2, \ldots, f_{n_{\text{nodes}}})$.

To simulate realistic sample purity, we define the node probabilities based on a mixture of healthy and cancerous cells. The sample purity, defined as the proportion of cancer cells in a sample and denoted by $\rho$, is drawn from a $\text{Beta}(9, 1)$ distribution, which places most of its mass in the range observed in real AML samples [1,3]. The probability of being assigned to the normal (non-cancerous) clone is set to $\pi_1 = 1 - \rho$, and the remaining probability mass $\rho$ is distributed across the cancer clones according to their relative frequencies at time point $s$. This yields the final node assignment probabilities at time point $s$:

$$(\pi_1, \pi_2, \ldots, \pi_{n_{\text{nodes}}}) = (1.0 - \rho, \rho f_2, \ldots, \rho f_{n_{\text{nodes}}}).$$

### A4 Simulations

#### A4.1 Longitudinal Simulation Details

Clonal frequencies were simulated using the Moran process described in Section A3. Figures A5–A8 display stacked plots of clonal dynamics for the 40 simulated cancers used to evaluate LoPhy and competing methods. Each color represents a distinct cancer clone population, with its vertical height indicating its proportion of the total cancer cell population at each time step during evolution. Grey dashed vertical lines indicate the sampling time points, labeled S1, S2, etc. Ten datasets were simulated for each simulation setting, with the dataset index shown in the top-left corner of each plot.

Figures A5–A8 demonstrate that our simulation framework generates diverse evolutionary trajectories, ranging from cases where a single clone dominates throughout evolution (e.g., Fig. A6, panel 9; Fig. A7, panel 6; Fig. A8, panel 6) to those where multiple clones expand and contract over time (e.g., Fig. A6, panel 4; Fig. A7, panel 9; Fig. A8, panel 2).

#### A4.2 Additional evaluations

We provide additional evaluations for LoPhy and competing methods on the 40 simulated longitudinal cancer datasets to supplement the results shown in Figure 2 of the main text. These supplementary analyses, presented in Figure A9, include precision, recall, and F1 scores for SNV–SNV pairs, CNA–CNA pairs, and all mutation pairs. For SNV–SNV pairs, we report only recall, as the number of SNVs is fixed between the inferred and ground truth trees. We also include runtime distributions for each method under each simulation setting in Figure A11. To further examine SNV–SNV recall, Figure A10 breaks down performance by the ancestral relationship between each SNV pair and reports the precision, recall, and F1 score relative to the ground-truth tree. The SNV pair relationships considered can be found in Section A2.

The results shown in Figure A9 and Figure A10 support that LoPhy, on average, produces the most accurate tree structures relative to the ground truth tree. These results also provide insight into how to improve the current version of LoPhy. Specifically, Figure A9 shows that both LoPhy and COMPASS struggle with accurate CNA inference, with errors arising from a combination of imprecise CNA calls and missed true events. Although COMPASS infers many erroneous CNAs—resulting in extremely low precision—its recall remains, on average, lower than LoPhy's. LoPhy's recall is still relatively limited, suggesting substantial room to improve the underlying modeling of total and mutant copy numbers.

#### A4.3 Single time point evaluation

We evaluated LoPhy on simulated single–sample cancer data to assess its applicability to both single–time point and longitudinal settings. We simulated 10 single–sample datasets, each comprising 20 genomic regions, 20 SNVs, 3 CNAs, and 6

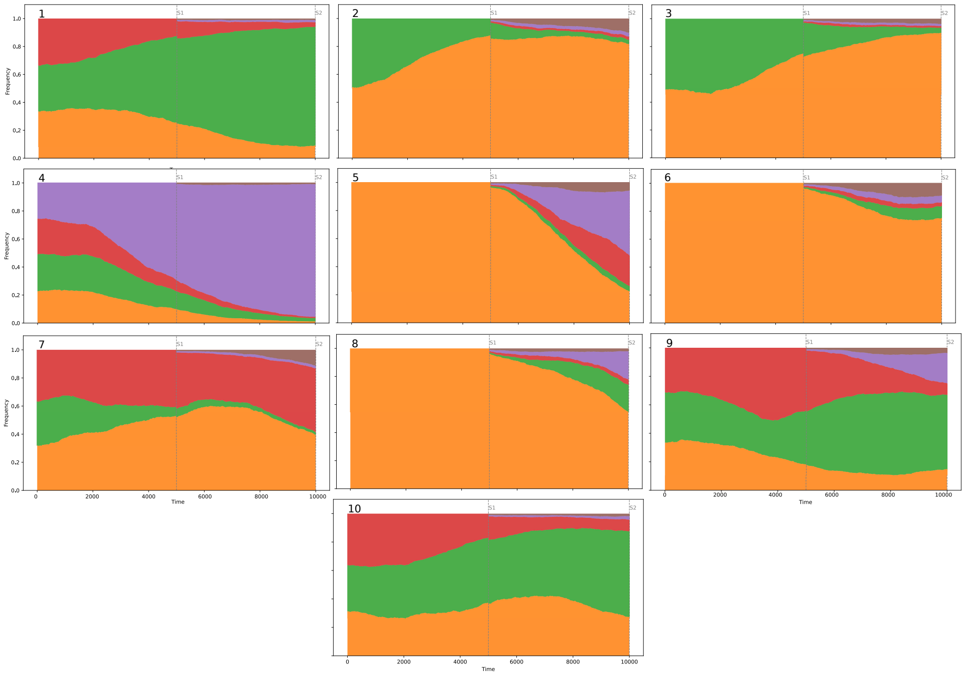


**Fig A5.** Clonal frequency dynamics for simulated cancers with two longitudinal samples.

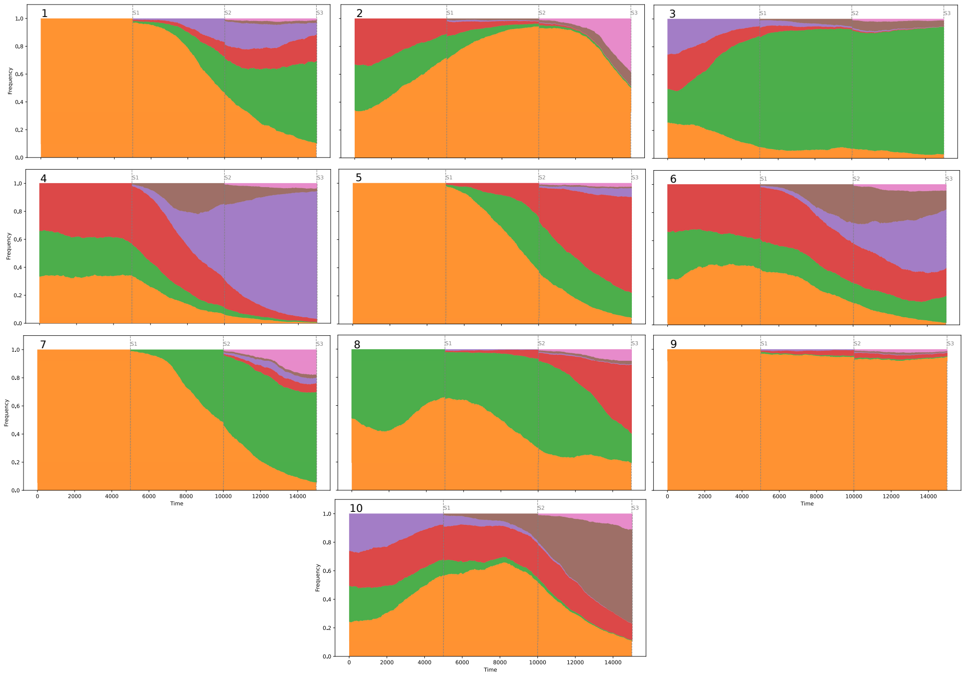


**Fig A6.** Clonal frequency dynamics for simulated cancers with three longitudinal samples.

clones. The results, shown in Figures A12 and A13, demonstrate that LoPhy can accurately reconstruct clone trees from single–time point data, often outperforming COMPASS and SCITE. While LoPhy achieves lower MCN-MAE and faster runtimes than COMPASS, its conservative CNA calling leads to slightly reduced CNA recall, resulting in marginally higher TCN-MAE and slightly lower Tree F1 scores. Because LoPhy's default parameters were not optimized for single–sample analyses, we tested reducing the penalty for introducing CNAs. This adjustment improves performance, yielding lower

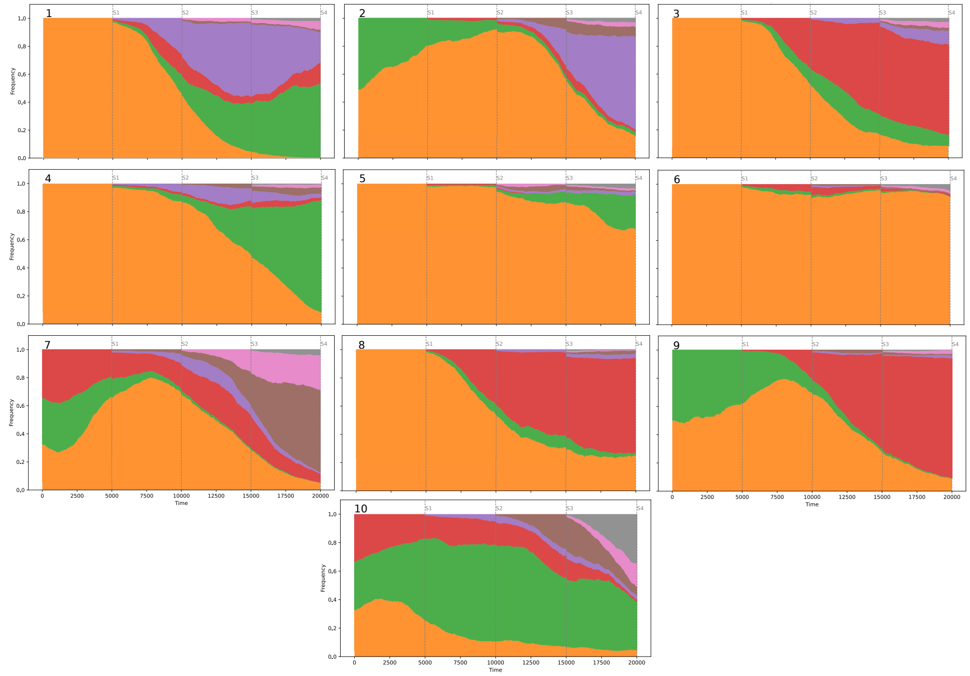


**Fig A7.** Clonal frequency dynamics for simulated cancers with four longitudinal samples.

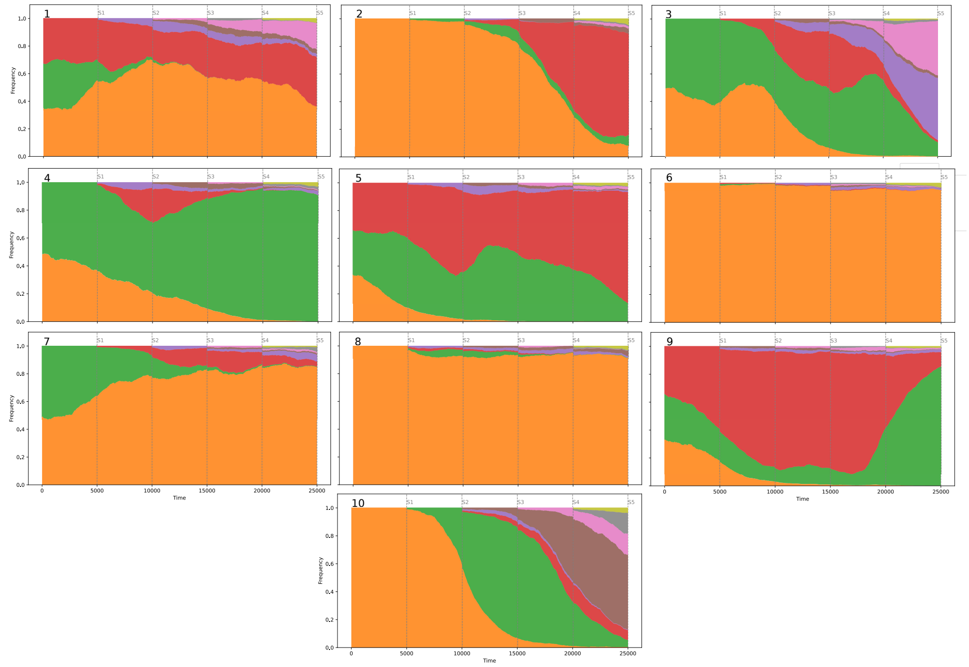


**Fig A8.** Clonal frequency dynamics for simulated cancers with five longitudinal samples.

TCN-MAE than COMPASS, perfect SNV–SNV and CNA–CNA F1 scores on all but one simulation, and overall Tree F1 scores only slightly below COMPASS (average difference 4e-4). Note that LACE was excluded from these analyses because it's specifically designed for longitudinal settings.

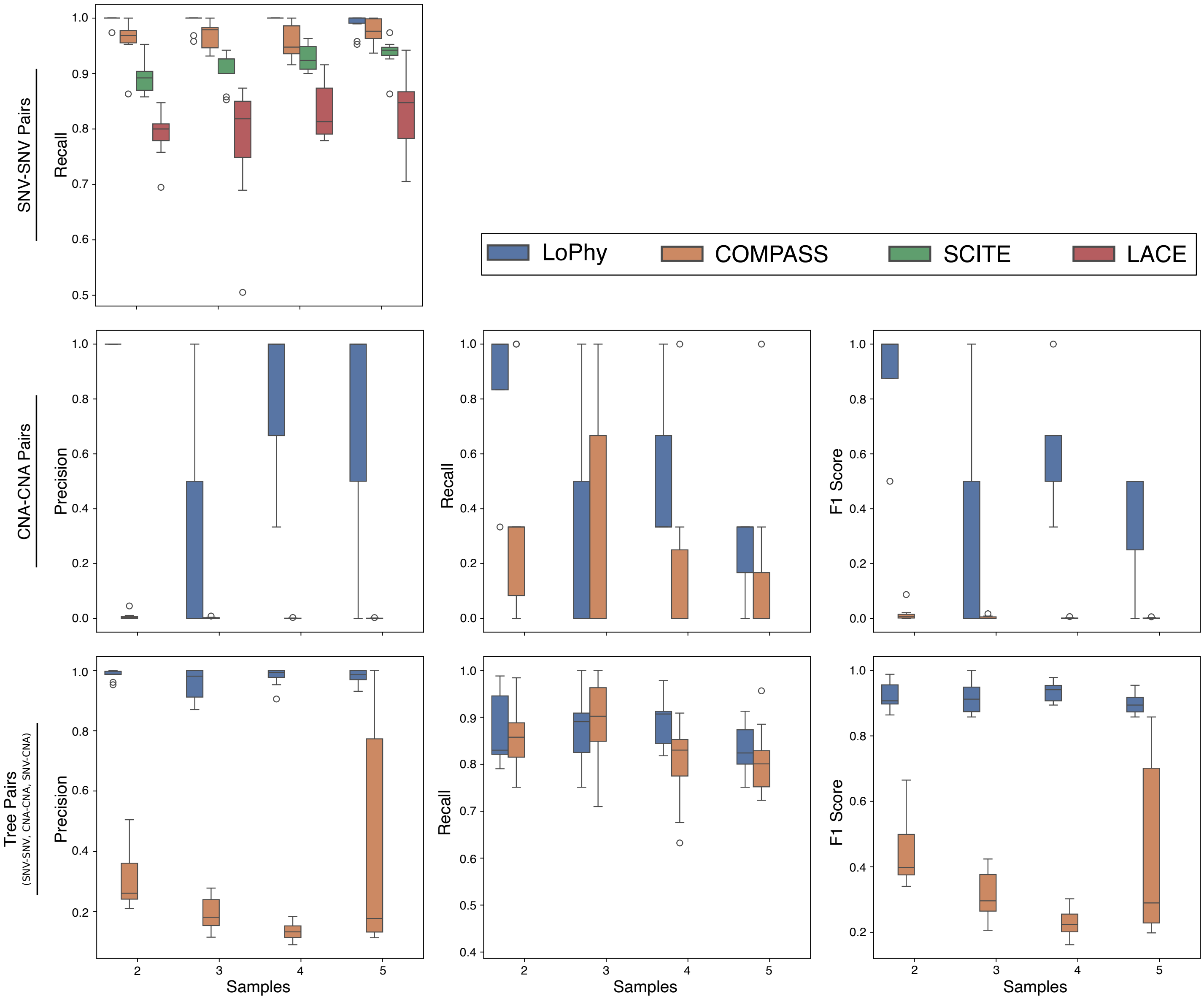


**Fig A9.** Additional evaluations of LoPhy, COMPASS, SCITE, and LACE on 40 simulated longitudinal cancer datasets. Boxplots show the distributions of precision, recall, and F1 scores for inferred mutation pairs (SNV–SNV, CNA–CNA, and SNV–CNA) relative to those in the ground-truth tree. CNA–CNA pair and Tree pair (all SNV and CNA combinations) metrics are shown only for LoPhy and COMPASS, as these are the only methods that infer CNAs.

### A5 Additional longitudinal AML reconstructions

The following sections compare and contrast reconstructions by LoPhy and COMPASS for the 15 AMLs from [1] and the 4 TP53-mutated AMLs from [2]. Note that AML-63 and AML-88 are excluded due to low data quality or extremely high sample purity.

Section 3.2 in the main text details the number of AML datasets where LoPhy or COMPASS inferred a longitudinally-observed clone tree inconsistent with the data or clinical description from [1]. The longitudinal data available for each AML are used to validate SNVs by examining single-cell variant allele frequencies (VAFs), and to validate CNAs by comparing region-level coverage patterns across groups of cells. The specific inconsistencies are categorized into three types: (1) false emergence—a mutation is inferred to occur earlier in evolution than indicated by the data or clinical description, (2)

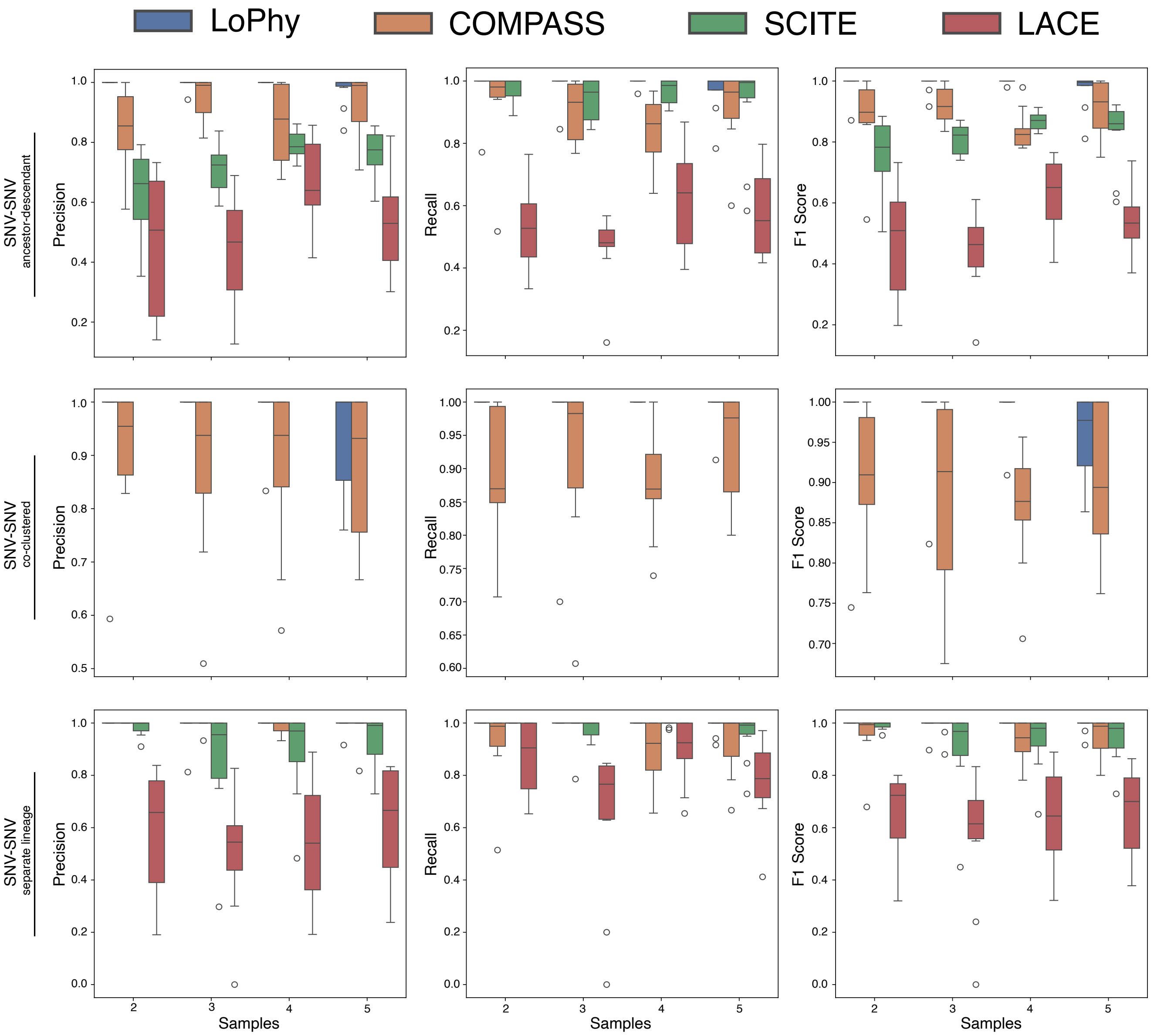


**Fig A10.** Evaluation of inferred SNV–SNV evolutionary relationships for LoPhy, COMPASS, SCITE, and LACE across 40 simulated longitudinal cancer datasets. Boxplots show the distributions of precision, recall, and F1 scores for each SNV–SNV ancestral relationship in the inferred trees, compared to the ground-truth trees. Only LoPhy and COMPASS are shown for the co-clustered SNV relationship, as these are the only methods that assign multiple SNVs to the same clone.

false persistence—a clone is inferred to persist in later samples despite being absent according to the data or clinical description, and (3) false contraction—a clone is inferred to contract in later samples despite evidence that it persisted. Using these definitions, LoPhy inferred a false emergence in AML-66. COMPASS inferred false emergences in AML-01, AML-09, and AML-21; false persistence in AML-04, AML-07, and AML-99; and a false contraction in AML-39.

#### A5.1 AML-01

Here, we further analyze the AML-01 trees reconstructed by LoPhy and COMPASS in Figure 4. Figure A14b provides additional support for LoPhy's FLT3 CNLOH placement: 85% of cells at diagnosis have a FLT3-ITD VAF $>65\%$, indicating the CNLOH event was likely present before the WT1 mutations. COMPASS infers two CNAs (gains on chr10 and chr16)

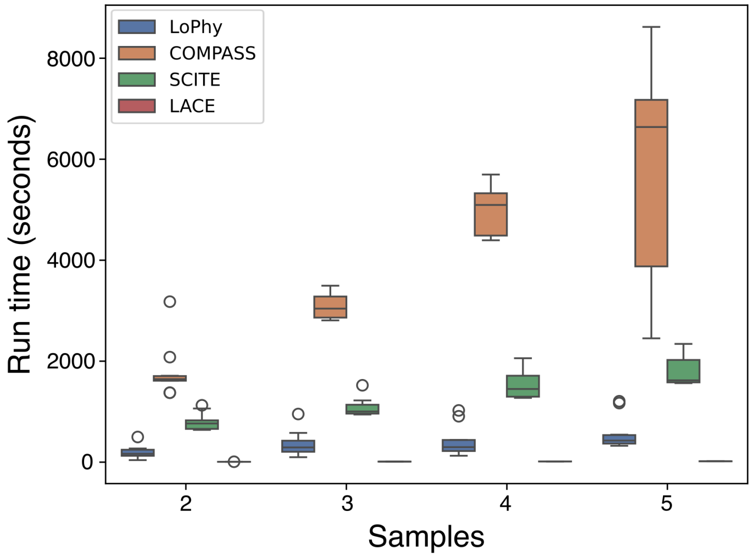


**Fig A11.** Run times for LoPhy, COMPASS, SCITE, and LACE on 40 simulated longitudinal cancers.

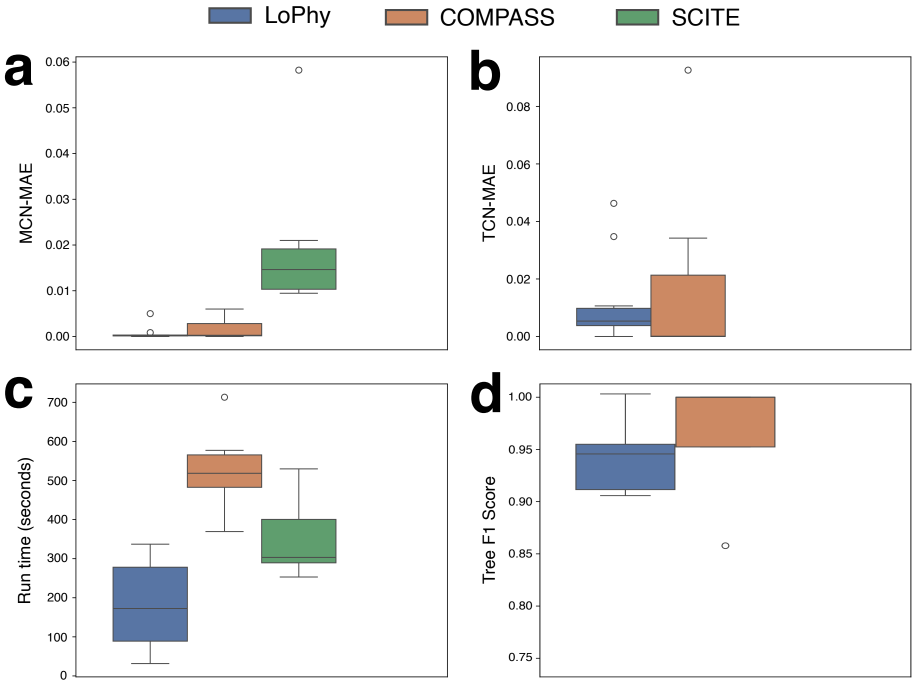


**Fig A12.** Evaluation of LoPhy, COMPASS, and SCITE on 10 simulated single–sample cancer datasets. **a–d.** Distributions of evaluation metrics, including MCN-MAE (*mutant copy number mean absolute error*), TCN-MAE (*total copy number mean absolute error*), Run time, and Tree F1 score.

coinciding with the NPM1 mutation. Figure A14c shows the fraction of reads mapped to chr10 and chr16 for 5,876 NPM1-positive cells at diagnosis versus 1,354 NPM1-negative cells. The relative difference in read fractions do not indicate gains, highlighting the pitfalls of COMPASS not modeling sample-specific variations in region coverage. No bulk sequencing data are available for AML-01 to confirm the FLT3 CNLOH event inferred by both methods.

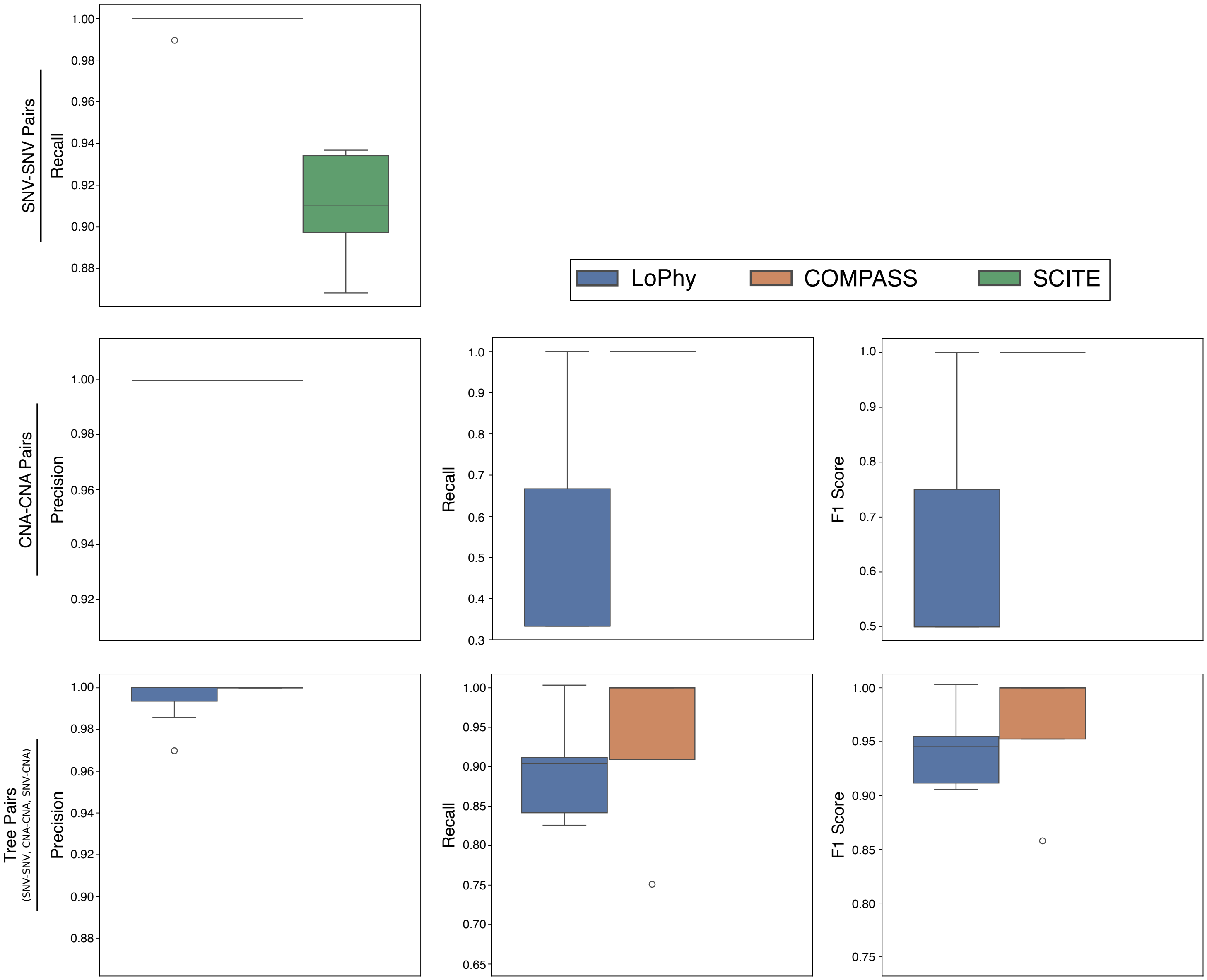


**Fig A13.** Additional evaluations of LoPhy, COMPASS, and SCITE on 10 simulated single-sample cancer datasets. Boxplots show the distributions of precision, recall, and F1 scores for inferred mutation pairs (SNV–SNV, CNA–CNA, and SNV–CNA) relative to those in the ground-truth tree. CNA–CNA pair and Tree pair (all SNV and CNA combinations) metrics are shown only for LoPhy and COMPASS, as these are the only methods that infer CNAs.

#### A5.2 AML-04

AML-04 was diagnosed with therapy-related FLT3-ITD AML (AML-04-001), with two additional samples collected after treatment: AML-04-002 (post azacitidine + quizartinib) and AML-04-003 (post crenolanib). Figure A15 shows the longitudinally-observed clone trees reconstructed by LoPhy and COMPASS, highlighting key differences: (1) COMPASS places the SF3B1 and SRSF2 mutations at the root of the tree (not shown) and detects no normal cells, and (2) LoPhy infers that the FLT3-ITD clone is present at diagnosis but lost after the first treatment, whereas COMPASS suggests the FLT3-ITD mutation remains in the dominant clones even after both treatments. The evolutionary history inferred by LoPhy aligns with clinical description in [1], which describe a dominant FLT3-ITD clone at diagnosis, followed by its decline and the emergence of a dominant SF3B1–SRSF2–IDH1 clone at later time points.

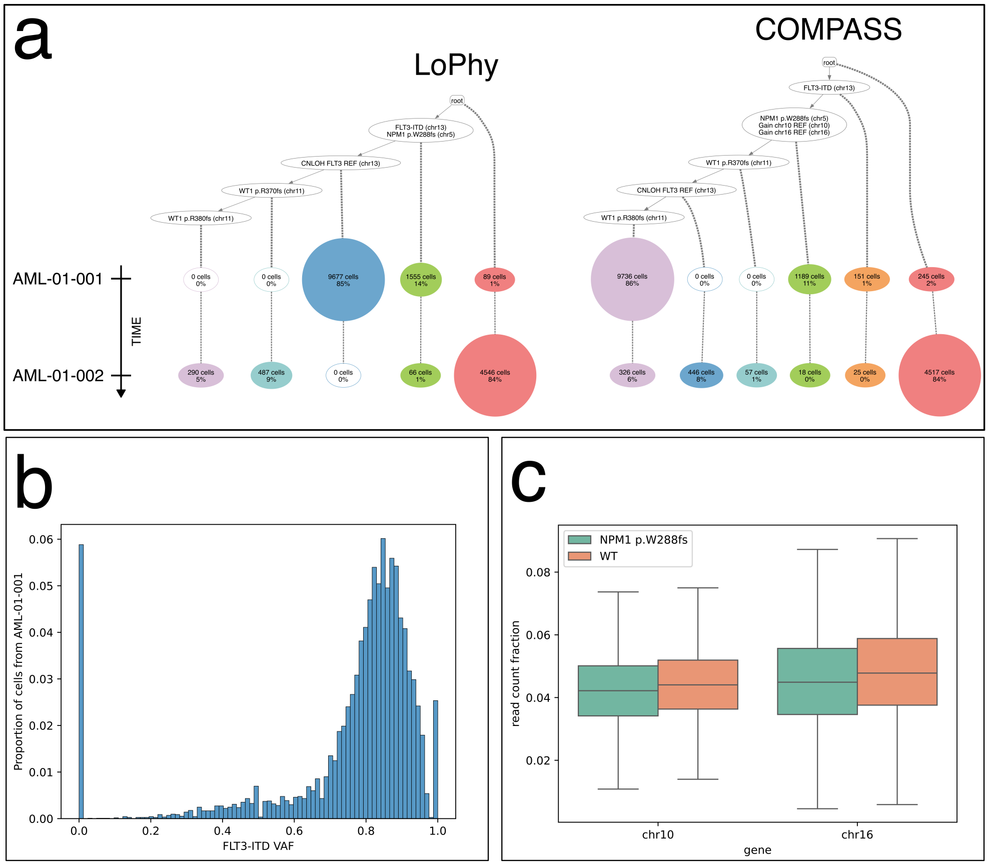


**Fig A14. a** Longitudinally-observed clone trees reconstructed by LoPhy and COMPASS for AML-01. **b.** Distribution of FLT3-ITD VAFs in cells at diagnosis (AML-01-001). **c.** Fraction of reads mapping to target regions chr10 and chr16 in sample AML-01-001 for NPM1-positive and NPM1-negative cells, indicating no copy number differences between the groups.

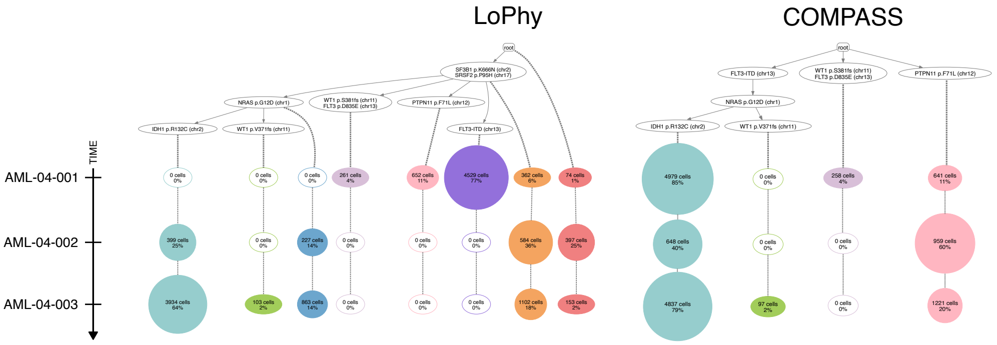


**Fig A15.** Longitudinally-observed clone trees reconstructed by LoPhy and COMPASS for AML-04.

#### A5.3 AML-07

AML-07 was diagnosed as pure erythroid leukemia (AML-07-001) with a RUNX1-mutated subclone, and later relapsed after treatment (AML-07-002). Figure A16 shows the longitudinally-observed clone trees reconstructed by LoPhy and COMPASS, which depict two markedly different evolutionary histories. LoPhy infers a dominant RUNX1 clone at diagnosis, characterized by a reference allele gain at FLT3 and loss of the reference alleles for RUNX1 and U2AF1. This clone shrank at relapse, where SRSF2 and IDH2 clones emerge as dominant. In contrast, COMPASS's tree suggests that the RUNX1 clone remains dominant at both time points, and is instead defined by two CNLOH events affecting the RUNX1 and U2AF1 reference

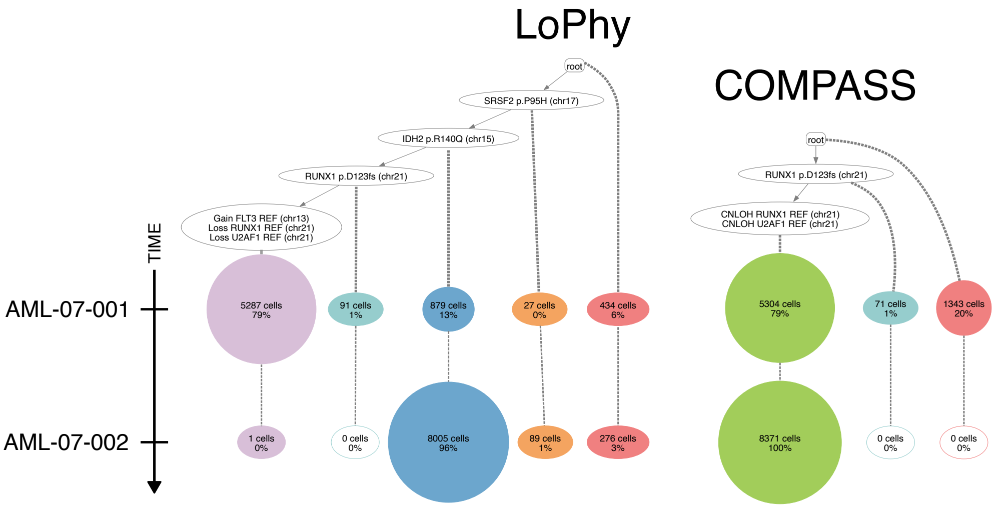


**Fig A16.** Longitudinally-observed clone trees reconstructed by LoPhy and COMPASS for AML-07.

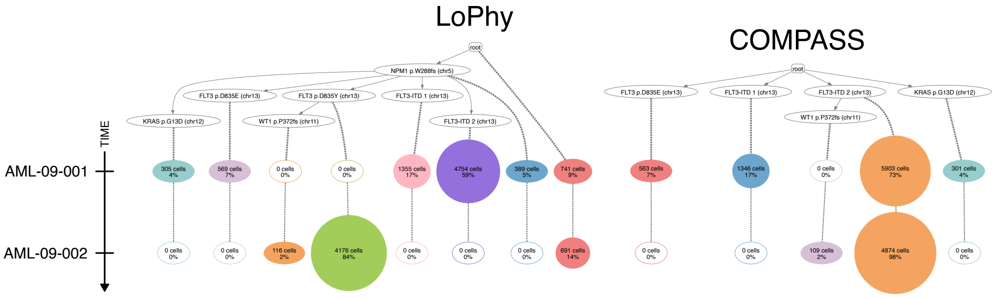


**Fig A17.** Longitudinally-observed clone trees reconstructed by LoPhy and COMPASS for AML-09.

alleles. LoPhy's reconstruction aligns with the clinical description in [1], which describe the RUNX1 clone as dominant at diagnosis but diminished at relapse, with IDH2- and SRSF2-characterized clones persisting at relapse. Notably, the authors in [3] used COMPASS to reconstruct a tree from only AML-07-001, and this single-timepoint tree closely agrees with LoPhy's longitudinal reconstruction. No bulk data is available for AML-07-001 to validate LoPhy's inferred CNAs.

#### A5.4 AML-09

AML-09 was diagnosed with therapy-related FLT3-ITD–mutated AML (sample AML-09-001), with a second sample collected after relapse (AML-09-002). Figure A17 shows the longitudinally-observed clone trees reconstructed by LoPhy and COMPASS. Both methods agree that numerous clones arose during evolution, characterized by distinct SNVs, but they differ substantially in the inferred clone proportions at the two time points. In LoPhy's tree, the FLT3-ITD 2 clone dominated at diagnosis but contracted by relapse, when the FLT3 p.D835Y clone expanded, became dominant, and subsequently acquired a WT1 mutation. This evolutionary history is consistent with the clinical description for this AML in [1]. In contrast, COMPASS places the FLT3 p.D835Y mutation at the root alongside NPM1. This occurs because COMPASS does not incorporate longitudinal information: seeing that most relapse cells carry FLT3 p.D835Y, it infers that all cells must harbor this variant, overlooking the absence of this mutation in the diagnosis sample.

#### A5.5 AML-18

AML-18 was diagnosed as AML with maturation (AML-18-001) and achieved remission following treatment, but later relapsed (AML-18-002). The clonal architecture was described as similar between diagnosis and relapse in [1]. Figure A18 shows the longitudinally-observed clone trees reconstructed by LoPhy and COMPASS. The main difference is that LoPhy separates the three SNVs into distinct clones, whereas COMPASS groups them into two clones and places IDH2 p.R172K at

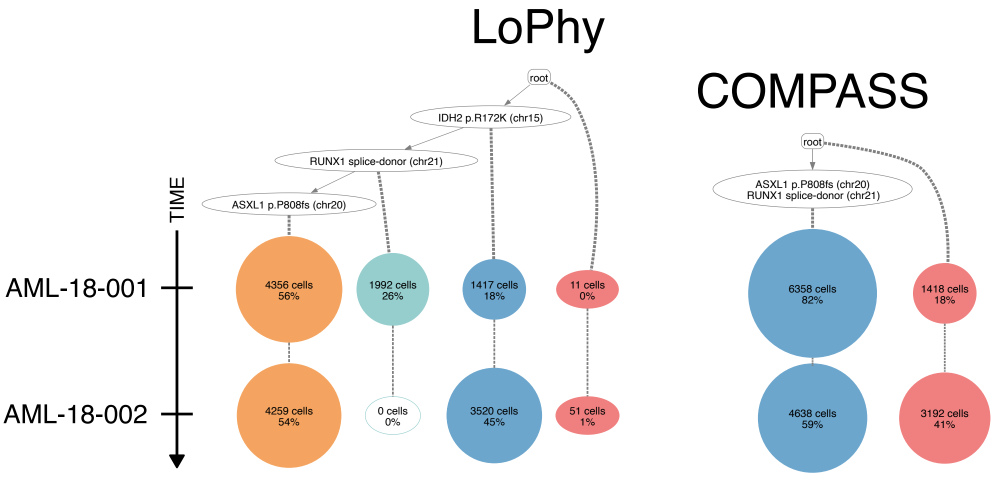


**Fig A18.** Longitudinally-observed clone trees reconstructed by LoPhy and COMPASS for AML-18.

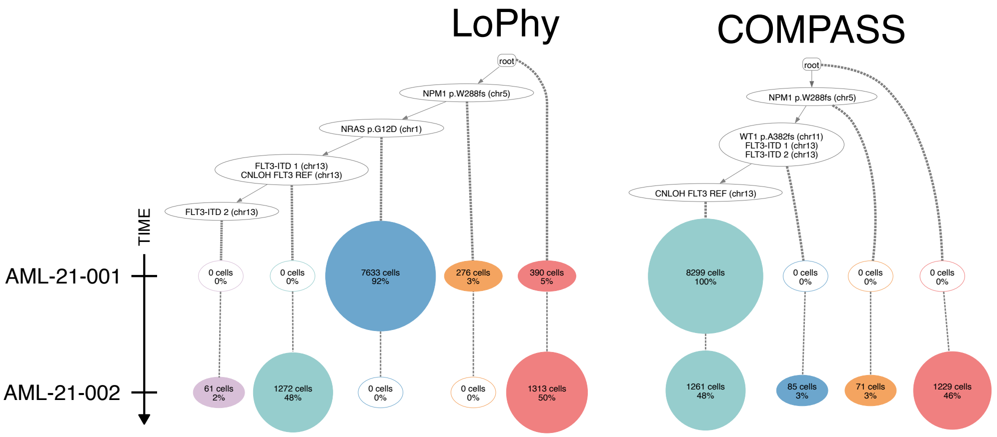


**Fig A19.** Longitudinally-observed clone trees reconstructed by LoPhy and COMPASS for AML-21.

the root. Both reconstructions are plausible, as the relapse sample (AML-18-002) shows high dropout of the RUNX1 splice-donor site, making it reasonable to infer that the RUNX1 mutation is either present or absent.

##### A5.6 AML-21

AML-21 had samples collected at diagnosis (AML-21-001) and relapse (AML-21-002). Figure A19 shows the longitudinally-observed clone trees reconstructed by LoPhy and COMPASS. Both methods identify a CNLOH event involving the FLT3 reference allele, but they differ in its inferred timing. LoPhy places the CNLOH in the same clone as the FLT3-ITD 1 mutation and before the acquisition of a second FLT3-ITD mutation. The FLT3 CNLOH-FLT3-ITD 1 clone is absent at diagnosis but becomes dominant at relapse. In contrast, COMPASS infers that the CNLOH occurs after both FLT3-ITD mutations, with this clone dominant at both diagnosis and relapse—an implausible result, since neither FLT3-ITD mutation was detected at diagnosis. The clinical description in [1] supports LoPhy's inference, reporting that the cancer was dominated by an NRAS clone at diagnosis, with WT1 and FLT3-ITD mutations undetected, and that at relapse the FLT3-ITD clone became dominant while the NRAS clone shrank.

##### A5.7 AML-38

AML-38 is a refractory AML with recurrent disease after multiple treatment cycles (AML-38-001, AML-38-002, AML-38-003). According to [1], the FLT3-ITD clone was largely eliminated after the first therapy cycle. Figure A20 shows clone trees reconstructed by LoPhy and COMPASS. Both reveal numerous subclones and infer CNLOH of the IDH2 alternative allele,

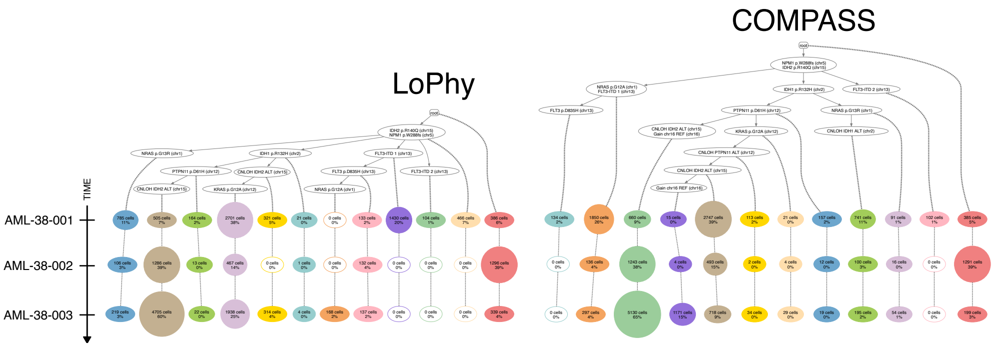


**Fig A20.** Longitudinally-observed clone trees reconstructed by LoPhy and COMPASS for AML-38.

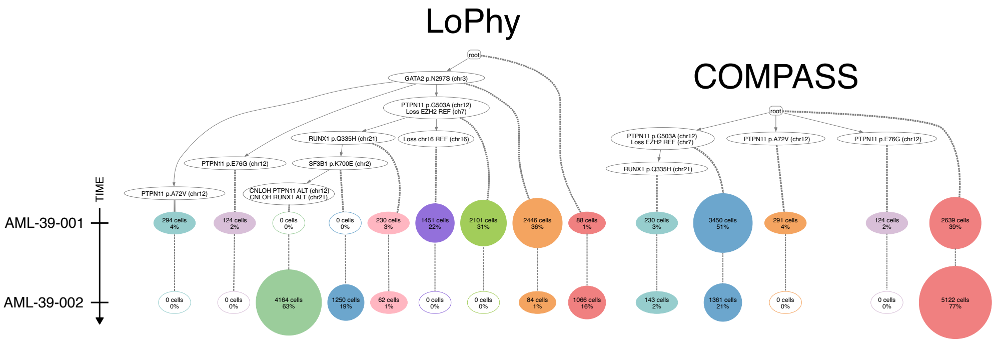


**Fig A21.** Longitudinally-observed clone trees reconstructed by LoPhy and COMPASS for AML-39.

though no bulk data is available for validation. They also agree that most FLT3-ITD–bearing clones disappear after the first cycle, while a few persist at low frequency in later samples. COMPASS additionally infers chromosome 16 gains, which cannot be validated.

#### A5.8 AML-39

AML-39 was diagnosed as a myelodysplastic syndrome that later progressed to secondary AML (AML-39-001). Complete remission was achieved after therapy, followed by relapse (AML-39-002). The clonal architecture was described as branching, with two clones harboring distinct PTPN11 mutations, both sharing SF3B1 and GATA2 mutations. At relapse, the PTPN11 p.A72V clone was eliminated, while the PTPN11 p.G503A clone was selected for and expanded.

Figure A21 compares longitudinally-observed clone trees reconstructed by LoPhy and COMPASS. LoPhy's tree closely matches the clinical description in [1], inferring that the two PTPN11 mutations formed independent branches and that the p.A72V clone, present at diagnosis, was lost at relapse, while the p.G503A clone and its descendants expanded to dominate. LoPhy further identifies CNAs within the PTPN11 p.G503A lineage, including EZH2 reference-allele loss (validated by bulk sequencing) and CNLOH affecting the alternative alleles of both PTPN11 and RUNX1 in the dominant relapse subclone. The single-cell VAFs in AML-39-002 support this structure: although $\sim$62% of cells exhibit VAFs $>$10% for the most recent mutation, SF3B1 p.K700E, these same cells show near-zero VAFs for the ancestral PTPN11 p.G503A mutation despite adequate coverage at that locus (median = 16, mean = 18), consistent with loss of the PTPN11 alternative allele via CNLOH. Likewise, these cells show near-zero VAFs for RUNX1 p.Q335H, with lower coverage (median = 0, mean = 8), supporting CNLOH of the RUNX1 alternative allele. Finally, the SF3B1 mutation mentioned clinically in [1] is not SF3B1 p.K700E, so the placement of this mutation in LoPhy's reconstruction remains consistent with the case description.

In contrast, COMPASS places most SNVs in the root clone, which it infers to be dominant at relapse, while the PTPN11 p.G503A clone contracts. COMPASS also infers EZH2 loss, but its tree matches exactly the reconstruction it produced when run on AML-39-001 alone [3].

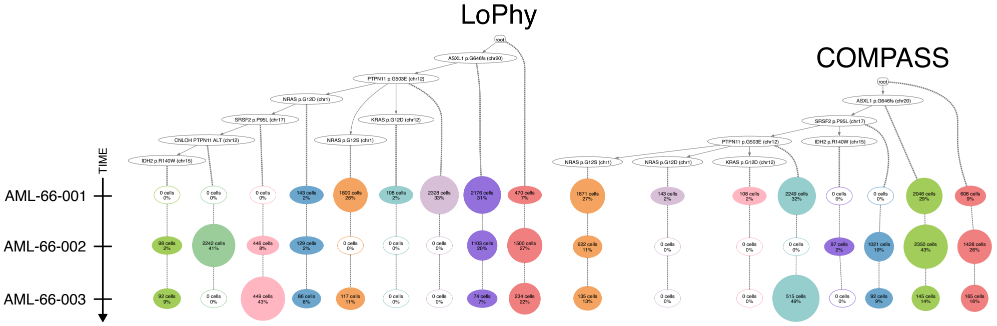


**Fig A22.** Longitudinally-observed clone trees reconstructed by LoPhy and COMPASS for AML-66.

#### A5.9 AML-66

AML-66 was sampled at diagnosis (AML-66-001), after treatment (AML-66-002), and at relapse (AML-66-003). Figure A22 compares longitudinally-observed clone trees reconstructed by LoPhy and COMPASS. Both depict branching evolution after PTPN11 p.G503E, giving rise to three subclones. LoPhy infers CNLOH of the PTPN11 alternative allele, which dominates AML-66-002. In this reconstruction, all cells in the IDH2 p.R140W clone carry this CNA, which is supported by the single-cell VAFs in AML-66-002. However, LoPhy assigns 92 cells from AML-66-003 to the IDH2 clone despite IDH2 p.R140W not being detected or adequately covered in the sample. With adequate IDH2 coverage, these cells would likely have been placed in the PTPN11 CNLOH clone (if IDH2 was absent) or a descendant clone (if IDH2 was present). These assignment errors may also reflect limited sampling, as AML-66-003 contained only 1,052 cells. In general, LoPhy is not designed to handle data with such sparse coverage between samples, as it expects mutations to accumulate over time and be covered in subsequent samples.

#### A5.10 AML-88

AML-88 was diagnosed as FLT3-ITD and FLT3 p.D835V positive AML (AML-88-001). Subsequent samples were collected after several therapy cycles (AML-88-002), at complete remission (AML-88-003), at relapse (AML-88-004), and during a second remission (AML-88-005). Unfortunately, due to missing data and high sample purity in the publicly available dataset, longitudinal reconstruction with LoPhy or COMPASS was not feasible.

#### A5.11 AML-99

AML-99 was sampled at diagnosis (AML-99-001), after multiple cycles of therapy (AML-99-002, AML-99-003), at relapse (AML-99-004), and following additional treatment (AML-99-005). LoPhy's longitudinal reconstruction is presented in Figure 3 of the main text and discussed in Section 3.2. Here, we present COMPASS's longitudinal reconstruction for AML-99 and provide a brief analysis.

Figure A23 shows the longitudinally-observed clone tree reconstructed by COMPASS for AML-99. While it correctly identifies a clone with RAD21 and MYC gains and clones defined by RUNX1 CNLOH, COMPASS incorrectly infers that several clones—including those carrying NRAS p.G60E and FLT3 p.K602fs—are present in all samples, despite these mutations appearing only in later samples according to [1]. Additionally, it infers SMC1A copy number changes and extensive branching for which there is limited supporting evidence.

#### A5.12 AML-107

AML-107 is a therapy-related AML, refractory to two treatments, with samples collected after each treatment (AML-107-001, AML-107-002). In [1], the clonal architecture was reported as unchanged across treatments. Figure A24b shows clone trees reconstructed by LoPhy and COMPASS, both depicting similar evolutionary histories with losses of FLT3, EZH2, and ETV6. Additionally, LoPhy infers a gain of the DNMT3A reference allele. The losses inferred by LoPhy are supported by coverage shifts in AML-107-001 between cells assigned to the root clone (baseline) and CNA-impacted clones (Figure A24a). Note that COMPASS places DNMT3A p.K202N in the root clone.

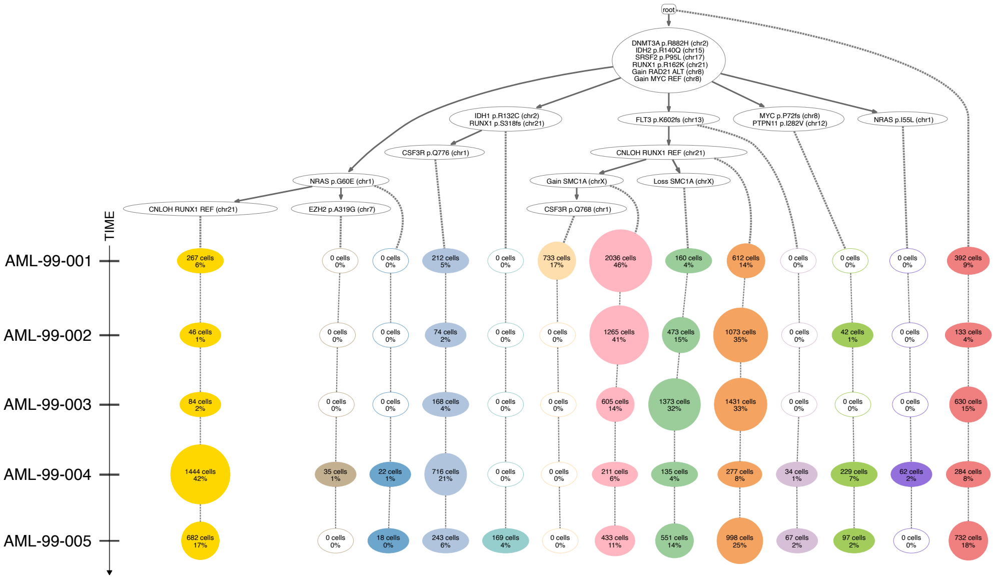


**Fig A23.** Longitudinally-observed clone tree reconstructed by COMPASS for AML-99.

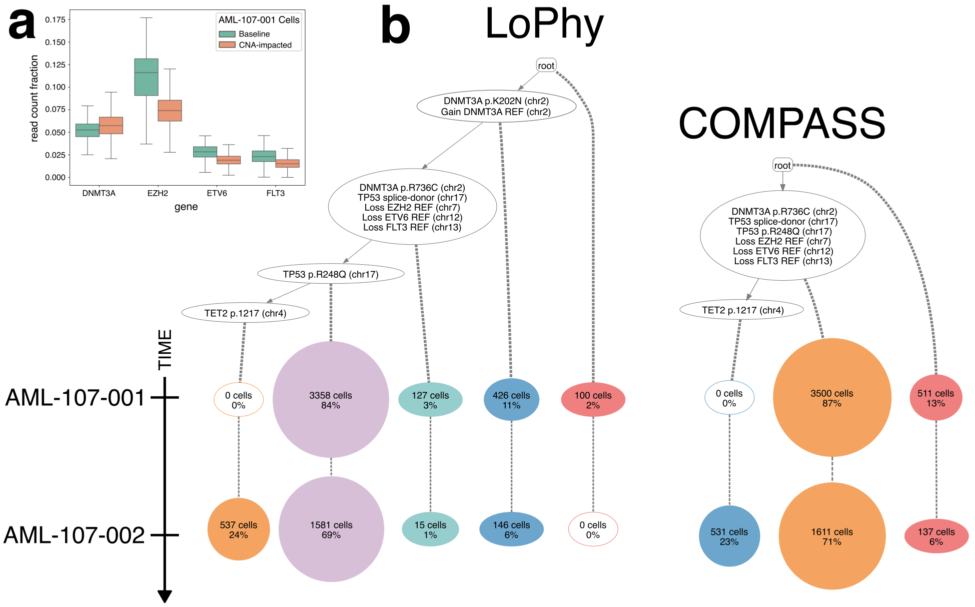


**Fig A24. a.** Coverage changes in AML-107-001 between cells assigned to the root clone (baseline) and those assigned to CNA-impacted clones. **b.** Longitudinally-observed clone trees reconstructed by LoPhy and COMPASS for AML-107.

#### A5.13 AML-CAL012

Figure A25 shows the longitudinally-observed clone trees for AML-CAL012 reconstructed by LoPhy and COMPASS. As reported in [2], the TP53 clone expanded from approximately 50% pre-treatment (AML-CAL012-001) to 80% post-treatment (AML-CAL012-002), while the PTPN11 and KRAS subclones grew from approximately 1% to 20%. Both reconstructions are consistent with this description.

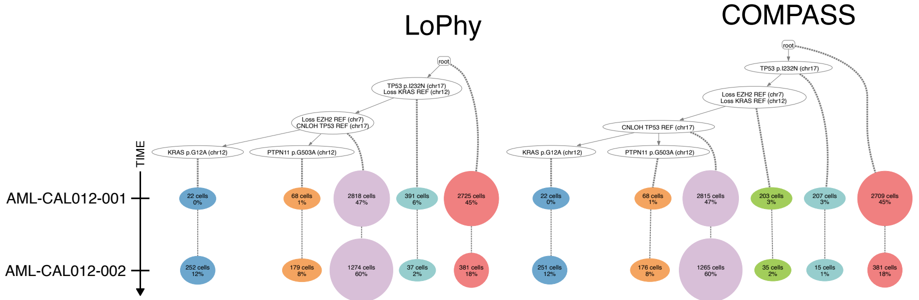


**Fig A25. a-b.** Longitudinally observed trees reconstructed by LoPhy and COMPASS for AML-CAL012.

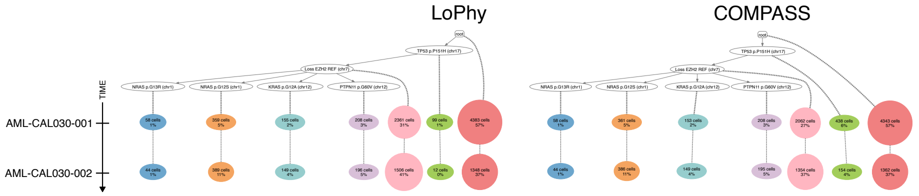


**Fig A26. a-b.** Longitudinally observed trees reconstructed by LoPhy and COMPASS for AML-CAL030.

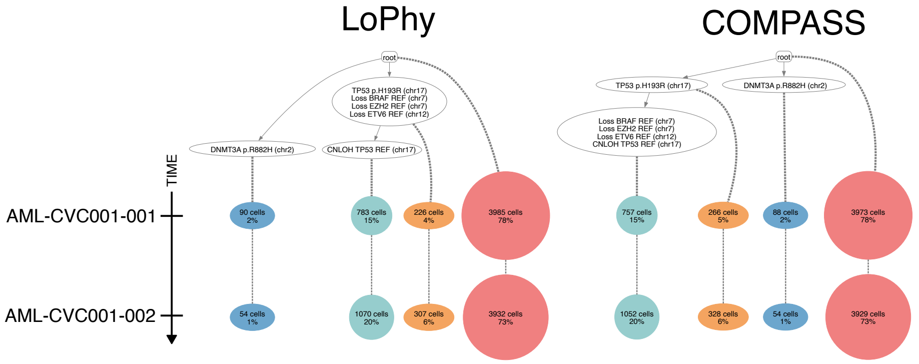


**Fig A27. a-b.** Longitudinally observed trees reconstructed by LoPhy and COMPASS for AML-CVC001.

#### A5.14 AML-CAL030

Figure A26 shows the longitudinally-observed clone trees for AML-CAL030 reconstructed by LoPhy and COMPASS. As reported in [2], the TP53 clone expanded from approximately 40% pre-treatment (AML-CAL030-001) to 60% post-treatment (AML-CAL030-002), while the PTPN11 and N/KRAS subclones grew from approximately 10% to 20%. Both reconstructions are consistent with this description.

#### A5.15 AML-CVC001

Figure A27 shows the longitudinally-observed clone trees for AML-CVC001 reconstructed by LoPhy and COMPASS. As reported in [2], the TP53 clone expanded modestly from approximately 20% pre-treatment (AML-CVC001-001) to 22% post-treatment (AML-CVC001-002), and both reconstructions capture this. Both trees include multiple CNAs, though these events cannot be validated due to the lack of bulk data for AML-CVC001.

#### A5.16 AML-CAU001

Figure A28 shows the longitudinally-observed clone trees for AML-CAU001 reconstructed by LoPhy and COMPASS. As reported in [2], the TP53 clone expanded from approximately 70% pre-treatment (AML-CAU001-001) to 80% post-treatment (AML-CAU001-002), and both reconstructions capture this expansion. Both trees include multiple CNAs, though these

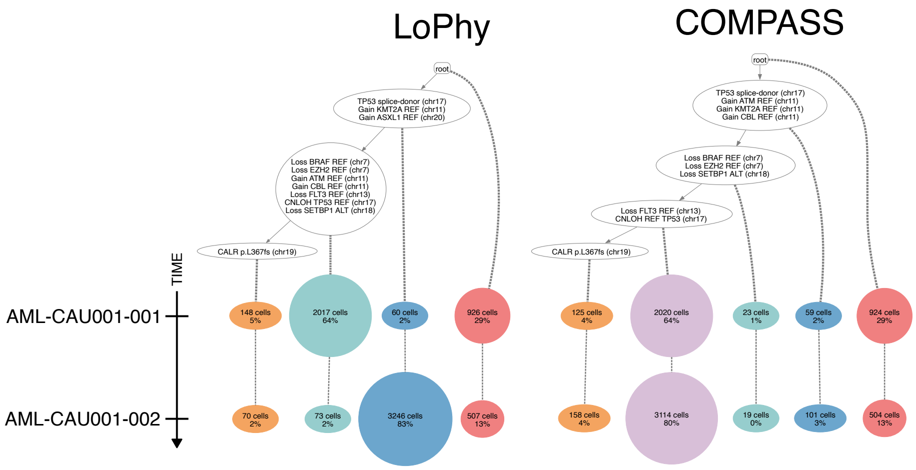


**Fig A28. a-b.** Longitudinally observed trees reconstructed by LoPhy and COMPASS for AML-CAU001.

cannot be validated due to the lack of bulk data for AML-CAU001. Overall, the reconstructions suggest a complex clonal architecture with numerous CNAs already present prior to treatment.
